# Automatic Ploidy Prediction and Quality Assessment of Human Blastocyst Using Time-Lapse Imaging

**DOI:** 10.1101/2023.08.31.555741

**Authors:** Suraj Rajendran, Matthew Brendel, Josue Barnes, Qiansheng Zhan, Jonas E. Malmsten, Pantelis Zisimopoulos, Alexandros Sigaras, Kwabena Ofori-Atta, Marcos Meseguer, Kathleen A Miller, David Hoffman, Zev Rosenwaks, Olivier Elemento, Nikica Zaninovic, Iman Hajirasouliha

## Abstract

Assessing fertilized human embryos is crucial for in vitro-fertilization (IVF), a task being revolutionized by artificial intelligence and deep learning. Existing models used for embryo quality assessment and chromosomal abnormality (ploidy) detection could be significantly improved by effectively utilizing time-lapse imaging to identify critical developmental time points for maximizing prediction accuracy. Addressing this, we developed and compared various embryo ploidy status prediction models across distinct embryo development stages. We present BELA (Blastocyst Evaluation Learning Algorithm), a state-of-the-art ploidy prediction model surpassing previous image- and video-based models, without necessitating subjective input from embryologists. BELA uses multitask learning to predict quality scores that are used downstream to predict ploidy status. By achieving an AUC of 0.76 for discriminating between euploidy and aneuploidy embryos on the Weill Cornell dataset, BELA matches the performance of models trained on embryologists’ manual scores. While not a replacement for preimplantation genetic testing for aneuploidy (PGT-A), BELA exemplifies how such models can streamline the embryo evaluation process, reducing time and effort required by embryologists.

## Introduction

Since the advent of in vitro fertilization (IVF) in 1978, it has served as a key solution for individuals unable to conceive naturally, accounting for over 8 million successful births globally.^1^ This procedure involves transvaginal transfer of laboratory-fertilized oocytes into the uterus. A critical determinant of IVF success and minimizing risk of perilous multiple pregnancies lies in the selection of high-quality, single normal embryos, primarily influenced by their ploidy status.^2,3^

Ploidy status, the chromosomal constitution of an embryo, greatly impacts pregnancy outcomes. Euploid embryos, characterized by normal chromosomal counts, typically lead to successful pregnancies, while aneuploid embryos—those with chromosomal aberrations—are associated with miscarriage, failed pregnancies, and chromosomal disorders like Down syndrome or Turner’s syndrome. Embryo aneuploidy, which leads to increased miscarriage rates, correlates with advanced maternal age.

Currently, preimplantation genetic testing for aneuploidy (PGT-A) is used to ascertain embryo ploidy status. This procedure requires a biopsy of trophectoderm (TE) cells, whole genome amplification of their DNA, and testing for chromosomal copy number variations. Despite enhancing the implantation rate by aiding the selection of euploid embryos, PGT-A presents several shortcomings.^4^ It is costly, time-consuming, and invasive, with potential to compromise embryo viability. Moreover, the test’s accuracy can be marred by embryonic mosaicism— the co-existence of aneuploid and euploid cells within the TE— leading to false results, diminished embryo viability, and lower implantation rates.^5^

The advent of computer vision in artificial intelligence, along with the accumulation of extensive IVF-related datasets—incorporating images, videos, and clinical outcomes—has spurred the development of automated embryo assessment methods via time-lapse image analysis. For instance, Khosravi et al. designed STORK, a model assessing embryo morphology and effectively predicting embryo quality aligned with successful birth outcomes.^6^ Analogous algorithms can be repurposed for embryo ploidy prediction, based on the premise that embryo images may exhibit patterns indicative of chromosomal abnormalities. Chavez-Badiola et al. employed the deep learning model ERICA to analyze 1,231 embryo images to predict ploidy status, achieving a 70% accuracy, with an area under the receiver operating characteristic curve (AUC) of 74%, and a sensitivity and specificity of 54% and 86% respectively. Notably, ERICA predicted a euploid embryo in the top rank in 79% of cases. Limitations included the model’s inability to distinguish between single and complex aneuploidies, the assumption of euploidy confirmation at β-HCG ≥20 mUI/mL on day 7, and a limited dataset potentially restricting general applicability.^7^ Similarly, Barnes et al. devised machine learning algorithms to predict embryo ploidy status from a single image at 110 hours post insemination (hpi), using time-lapse sequences.^8^ Silver et al. speculated that the entirety of video sequences could potentially improve embryo classification accuracy, leading to the development of the UBar CNN-LSTM model, which attained an AUC of 0.82—though on a limited dataset.^9^ In another recent study, Lee et al. utilized a two-stream inflated 3D model on 670 image sequences, achieving an AUC of 0.74 in differentiating euploid/mosaic and aneuploid embryos.^10^

Analyzing entire time-lapse sequences of embryo development presents a challenge in predicting ploidy status, as not all developmental stages may provide pertinent information. This has led to previous studies focusing on feature extraction from specific developmental periods.^11^ Campbell et al. proposed the timing and presence of blastocyst expansion on day 5 as a predictor of ploidy status.^12^ However, this criterion’s predictive accuracy has exhibited considerable variability across clinics, making it less reliable.^13^ Analyzing full embryo development videos could bypass the need to pinpoint relevant timeframes, but the computational cost of training models on vast datasets could compromise performance due to noise. Addressing these challenges, we present BELA—a fully automated ploidy prediction model—that requires only embryo time-lapse sequences and maternal age as inputs. By removing the need for subjective manual annotation, BELA not only streamlines the ploidy prediction process but also fosters broad applicability across different clinical settings.

## Results

### Training and Validation Datasets

In our study, we utilized deep learning techniques to predict ploidy status using time-lapse sequences of embryo development and compared various model performances across multiple clinics. Two internal datasets from Weill Cornell Medicine’s Center for Reproductive Medicine (WCM) were employed: the first encompassed 1,998 Embryoscope® time-lapse sequences, and the second contained 841 sequences from the Embryoscope+®. These sequences typically constituted 360-420 distinct frames, captured at 0.3-hour intervals over five days of development. PGT-A results served as the ground truth for ploidy prediction tasks, with embryos classified as euploid (EUP) or aneuploid (ANU). Further categorization of ANU embryos identified single aneuploid (SA)—with one chromosomal abnormality—and complex aneuploid (CxA)—with multiple chromosomal abnormalities. Accompanying clinical information included blastocyst scores (BS)—derived from morphological grades and morphokinetic parameters—and maternal age at oocyte retrieval. BS encompasses three sub-components: inner cell mass (ICM), trophectoderm (TE), and expansion score.^14^ This blastocyst score formulation has been shown to be predictive of implantation success, euploidy, and live-birth.^14^ For additional model validation, we utilized an external dataset from IVI Valencia, Spain. Unlike the WCM datasets, this dataset only contained EUP/ANU labels without explicit SA/CxA details and BS. A second external dataset from IVF Florida provided additional detail allowing discrimination between SA and CxA embryos. Comprehensive descriptions of these datasets are detailed in **Table 1, Supplemental Table 10**, and further expounded in the Methods section.

**Table 1.** Characteristics of Datasets. The sample size, distribution of data across ploidy classes, and additional clinical features for each dataset are shown.

| Dataset | WCM-Embryoscope | WCM-Embryoscope+ | Spain | Florida |
| --- | --- | --- | --- | --- |
| Sample Size | 1,998 | 841 | 543 | 869 |
| Ploidy Splits | SA: 494<br>CxA: 588<br>EUP: 916 | SA: 170<br>CxA: 261<br>EUP: 410 | ANU: 309<br>EUP: 234 | SA: 202<br>CxA: 222<br>EUP: 445 |
| Clinical Features | Maternal Age<br>Blastocyst Score<br>ICM Score<br>TE Score<br>Expansion Score | Maternal Age<br>Blastocyst Score<br>ICM Score<br>TE Score<br>Expansion Score | Maternal Age | Maternal Age<br>Blastocyst Score<br>ICM Score<br>TE Score<br>Expansion Score |

### Ploidy Prediction Model with Model-derived Blastocyst Score

We introduce BELA, the Blastocyst Evaluation Learning Algorithm for ploidy prediction, a fully automated model detailed in **Figure 1**. The model comprises two steps. First, BELA predicts the Blastocyst Score (BS) from processed day-5 time-lapse videos (96 -112 hpi), a timeframe chosen based on our ablation analyses comparing embryonic development timepoints and image versus video inputs (**Supplemental Text 1**). The input video undergoes processing and transformation into feature vectors via a pre-trained spatial feature extraction model (**Figure 1**, steps 1 - 4). To optimize performance, we used a multitasking BiLSTM model to concurrently predict ICM, TE, expansion, and blastocyst score. We evaluated the first component of BELA using the mean absolute error (MAE). In the second step, BELA uses the now ‘model-derived blastocyst score’ (MDBS) to predict the embryo’s ploidy status, employing a logistic regression that integrates maternal age as a continuous input feature, as illustrated in **Figure 1**. We trained and evaluated BELA on EUP versus CxA and EUP versus ANU splits. BELA was trained on data from the WCM-Embryoscope dataset via four-fold cross-validation. Performance was gauged using accuracy, AUC, precision, and recall across the datasets from WCM-Embryoscope, WCM-Embryoscope+, Spain, and Florida. For comparison, we trained two baseline models using the same cross-validation splits. The first baseline is a day-5 video model which exclusively uses time-lapse input from 96 - 112 hpi to directly predict ploidy status using a BiLSTM architecture. The second baseline is an embryologist-annotated model which uses only the ground-truth BS to predict ploidy status using a logistic regression.

**Figure 1.**
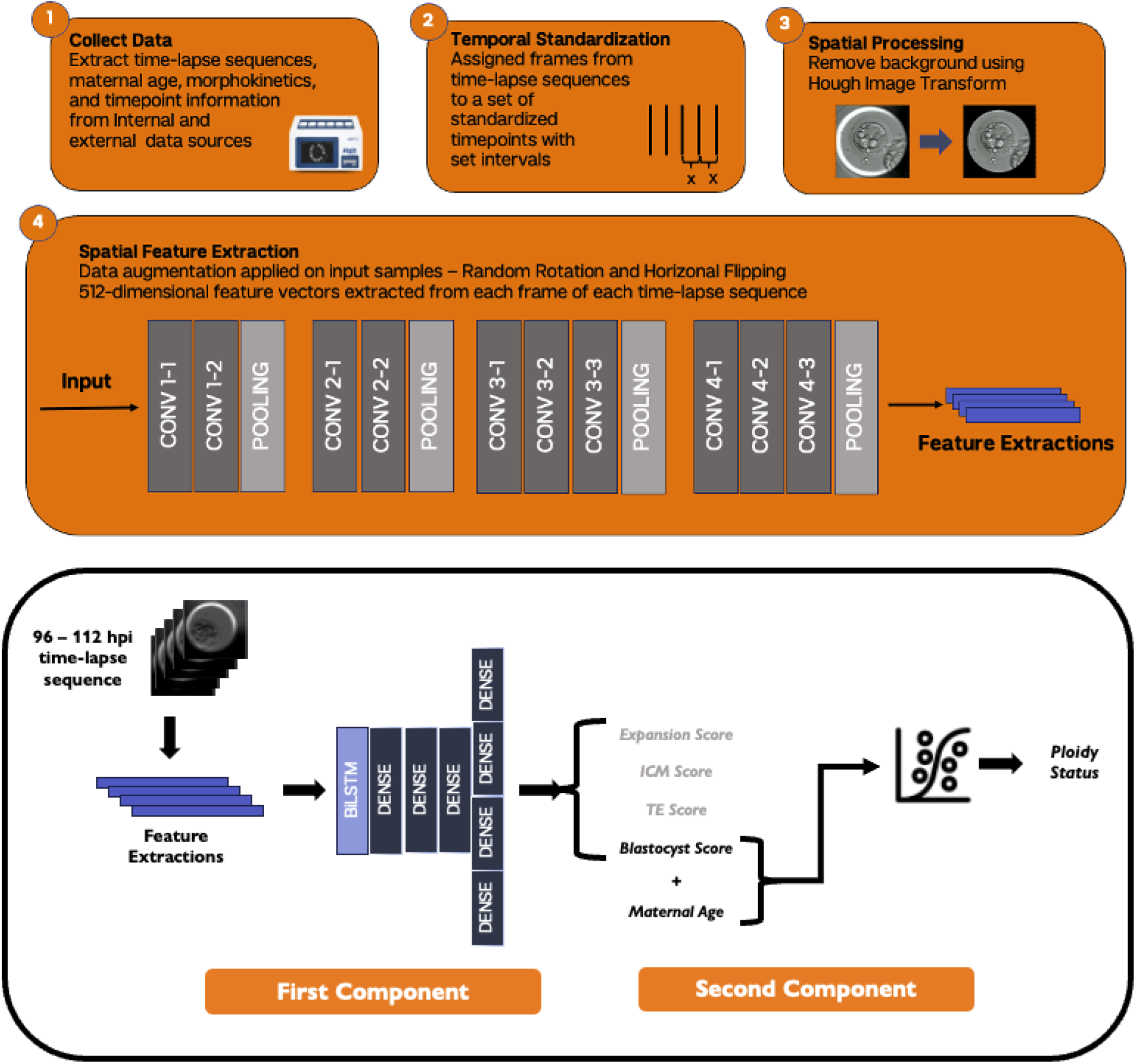
Overview of BELA development. Features are extracted from time-lapse image frames as shown in steps 1-4. Time-lapse images are both temporally and spatially processed to decrease bias. Horizontal and rotational augmentation is performed on time-lapse sequences. 512-dimensional features are extracted for each time-lapse image using a pretrained VGG16 architecture. These features are fed into a multitask BiLSTM model which is trained to predict blastocyst score as well as other embryologist-annotated morphological scores. Predicted blastocyst scores are inputted into a logistic regression model to perform ploidy prediction.

The first component of BELA predicts the blastocyst score (BS). As depicted in **Supplemental Figure 1**, both the training and test sets from WCM-Embryoscope show a moderate correlation (Pearson correlation >0.7) between the model-derived blastocyst scores (MDBS) and the embryologist BS. This moderate correlation is also evident in the predicted and actual scores of other embryologist metrics (**Supplemental Figure 2**). The mean absolute error (MAE) between the MDBS and the ground truth Embryoscope BS scores is 1.855 ± 0.03. The second phase of BELA involves ploidy classification. Using the WCM-Embryoscope test set, BELA, when trained to distinguish between EUP and ANU, attained an AUC of 0.66 ± 0.008, which rose to 0.76 ± 0.002 upon inclusion of maternal age. In the EUP versus CxA task, the AUC of the model was 0.708 ± 0.004 and increased to 0.826 ± 0.004 with the inclusion of maternal age. Comprehensive performance metrics of BELA are found in **Supplemental Table 1**. BELA’s performance (in orange), compared with a day-5 Video model and the embryologist-annotated blastocyst score model, is illustrated in **Figure 2**. In all tested scenarios (including or excluding age), test sets, and prediction tasks (EUP versus ANU and EUP versus CxA), BELA outperforms the day-5 video model (p < 0.05). Without including maternal age in ploidy prediction, the embryologist-annotated BS model surpasses BELA (p < 0.05) in all prediction tasks, barring EUP vs ANU on the WCM-Embryoscope test set (**Figure 2a**). However, with maternal age incorporated, BELA outperforms the embryologist-annotated blastocyst score model on the WCM-Embryoscope test set (p < 0.05). Still, it underperforms in comparison to the embryologist-annotated blastocyst score model on the WCM-Embryoscope+ dataset.

**Figure 2.**
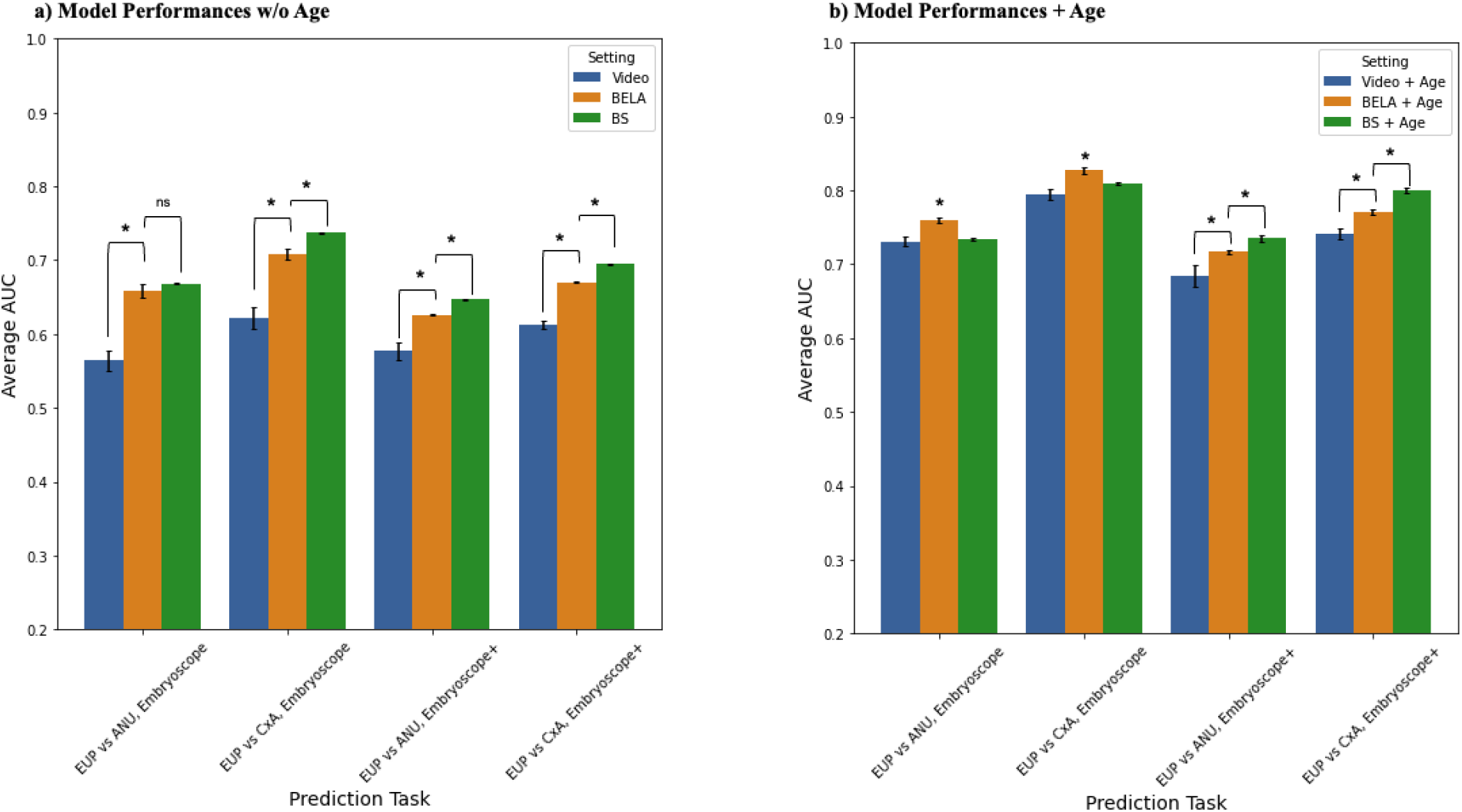
Comparison of BELA models with other models. Mean AUC scores and standard deviation for day-5 video, BELA, and embryologist-annotated BS trained models are shown. Performances are shown on both the WCM-Embryoscope and WCM-Embryoscope+ dataset for both prediction tasks. (a) Performances of models without maternal age. (b) Performances of models with maternal age.

The performance of the BELA was compared with a day-5 video model using an external dataset from Spain, consisting of 543 embryos (**Figure 3, Supplemental Table 1**). As the Spanish dataset includes only embryos labeled as ANU or EUP, model performance could only be measured for the task of distinguishing between EUP and ANU. Notably, BELA significantly outperforms the day-5 video model in both scenarios – with and without the inclusion of maternal age (p < 0.05). Unlike the embryos from Weill Cornell Medicine (WCM-Embryoscope and WCM-Embryoscope+ datasets), those from the Spanish dataset were artificially hatched on day 3, which likely impacted later blastocyst morphology and morphokinetics. These embryos exhibit bleached zona pellucida and lack the full expansion seen in the embryos from the training set. To quantitatively verify these differences, feature encodings were extracted using the pre-trained feature extractor for each frame (between 96 hpi and 112 hpi) of each embryo. Averaging these feature encodings across frames yielded a single feature encoding for each embryo, which was further dimensionally reduced via PCA. The resulting feature encodings, categorized by dataset, can be viewed in **Supplemental Figure 3**. The feature space shows a significant overlap between the datasets based in the United States, while the Spanish data clusters distinctly towards the bottom right. However, despite these noticeable differences, the performance of the model (excluding maternal age) remains comparable to that achieved with the Weill Cornell Medicine datasets. This suggests that the model might be generally applicable across various clinics, even those with practices that the training data did not account for. The models incorporating maternal age showed decreased performance relative to the Weill Cornell datasets, likely attributable to demographic differences among patients using IVF between Weill Cornell and Spain. For example, in Spain, IVF is more affordable and accessible due to different healthcare insurance policies, whereas in the United States, the high cost of IVF can limit its accessibility to individuals with the necessary financial resources.^15,16^ This likely contributes to the different maternal age distributions observed within the datasets.

**Figure 3.**
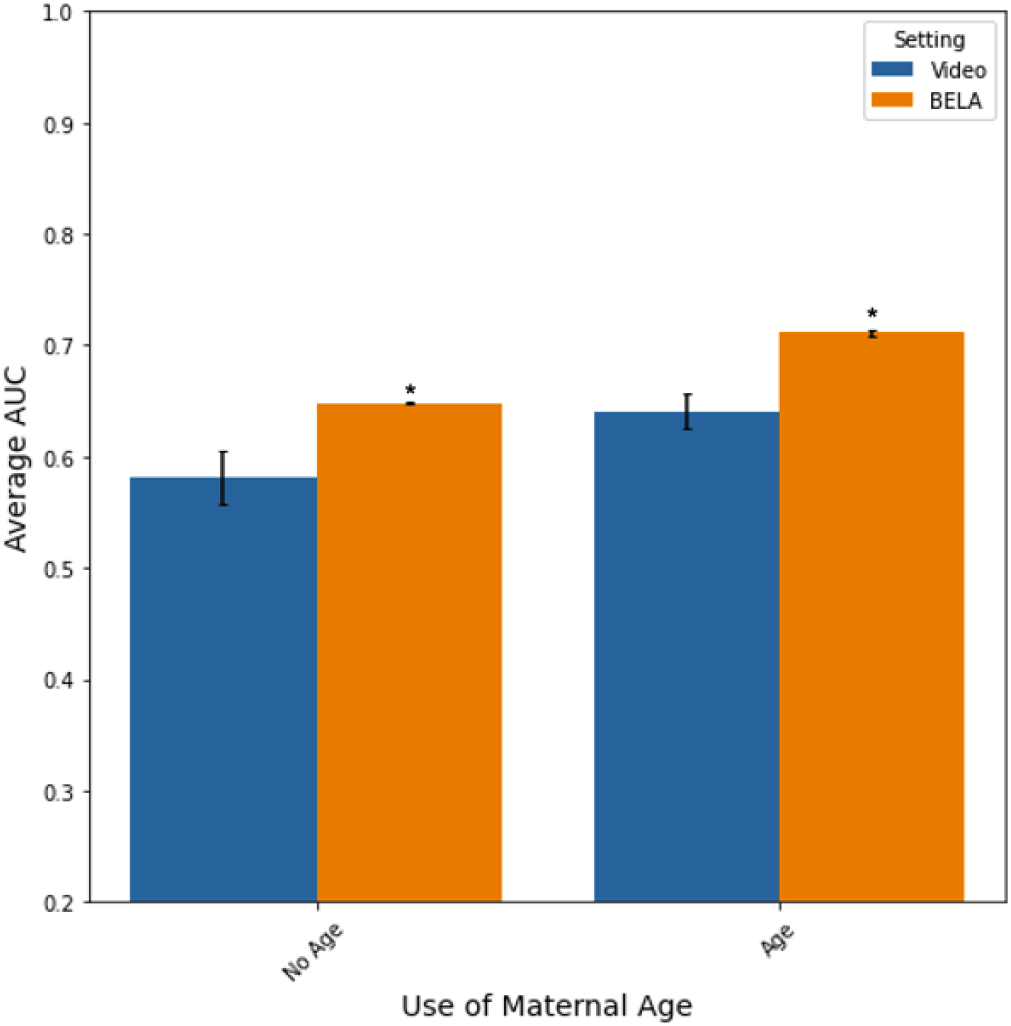
Performances of day-5 video model and BELA on the Spain dataset. Average AUC with standard errors is shown for all EUP vs ANU prediction tasks on the Spain dataset for both the day-5 video model and BELA. Blue bars depict model performances of day-5 video models, whereas orange bars depict performances of BELA models.

The performance of BELA was further assessed using an external dataset from IVF Florida, comprising 869 embryos. It is important to note that the performance of BELA, as well as comparison logistic regression models trained only on maternal age and embryologist-derived blastocyst score, significantly declined in comparison to the other test sets. This decrease in performance could be attributed to the weak correlations between blastocyst score, maternal age, and ploidy within the Florida dataset (**Supplemental Table 8**). Maternal age is a crucial predictor of ploidy in all our models, thus, any decrease in its correlation to ploidy can significantly impact performance. Moreover, the embryologist-derived blastocyst scores in the Florida dataset were predominantly centered around a score of 7, thereby reducing the granularity that made it a potent predictor of ploidy in other datasets. This lack of granularity within the Florida dataset might be a result of different scoring practices, as IVF Florida evaluates blastocysts at 115 and 144 hpi, in contrast with the methods employed at Weill Cornell, which also utilize earlier time points for determining blastocyst score. Interestingly, the model-derived blastocyst score (MDBS) from the first module of BELA shows a stronger correlation (−0.119) with ploidy status than the embryologist-derived blastocyst score (−0.101). This finding suggests that BELA can create a score mapping that aligns better with ploidy status compared to the original embryologist-derived blastocyst scores. In order to further validate this hypothesis, an embryologist at Weill Cornell re-graded the 50 embryos within the Florida dataset where the MDBS deviated most significantly from the provided Florida blastocyst scores. The scoring method at Weill Cornell allows for greater granularity in assessing embryo quality. We observed a decrease in mean absolute error (MAE) between the MDBS versus the re-graded blastocyst score from Weill Cornell (4.16) and the MDBS versus the original Florida blastocyst score (5.02). This suggests a higher agreement between the MDBS and the Weill Cornell scoring method. This improved mapping could explain why BELA, with maternal age included, significantly outperforms the model trained on maternal age and embryologist-derived blastocyst score for the EUP vs ANU task (p < 0.05) (**Figure 4**).

**Figure 4.**
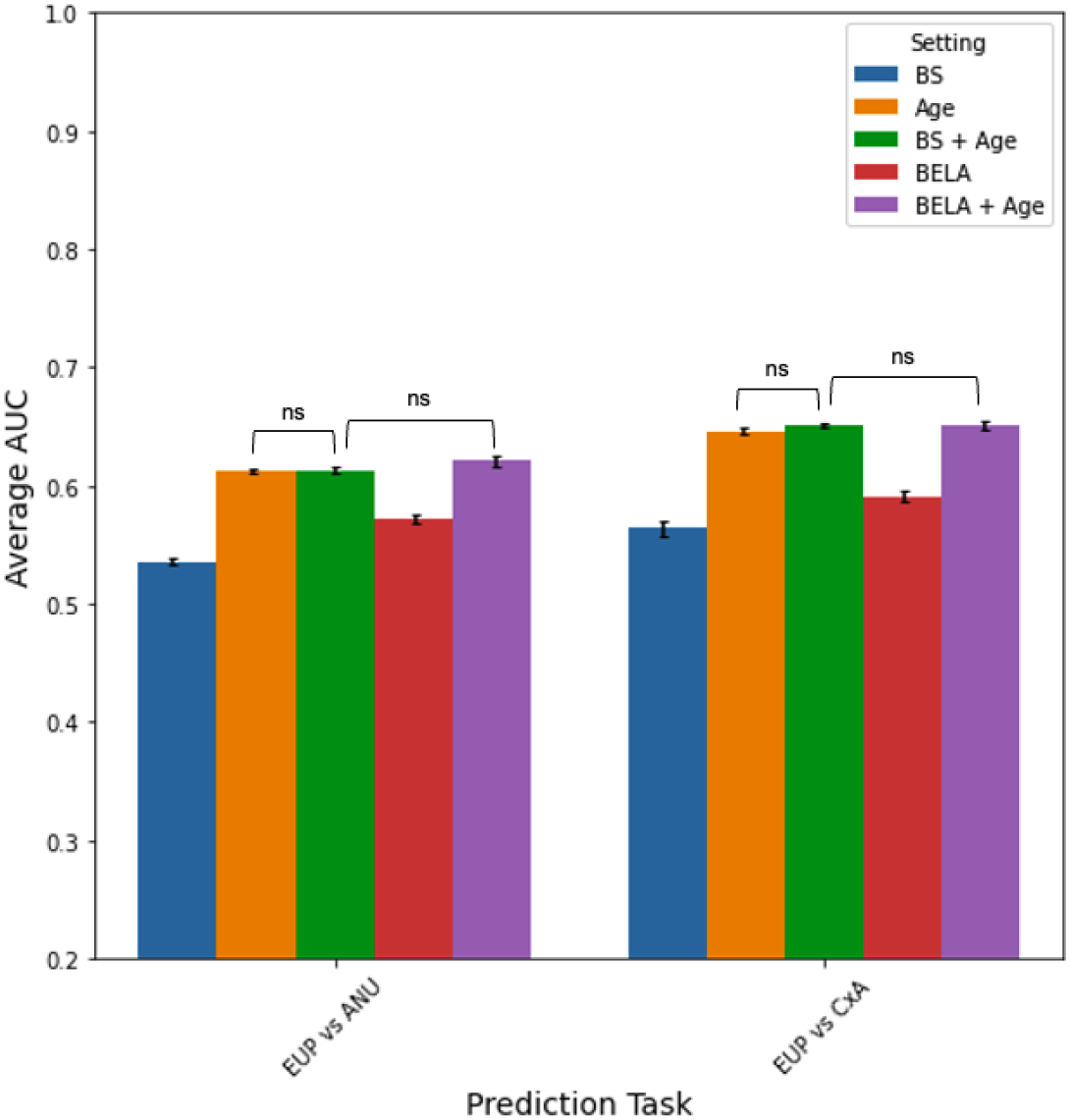
Performances of BELA and traditional machine learning models on the Florida dataset. Average AUC with standard errors is shown for EUP versus ANU and EUP versus CxA prediction tasks on the Florida dataset for BELA and logistic regression models trained only on embryologist-derived BS and/or maternal age.

In order to make the model available for clinical use, a web-based application named STORK-V for BELA was developed (**Figure 5, Supplemental Figure 4**). This platform is designed to be user-friendly and is capable of predicting an embryo’s ploidy status. The required input for the prediction includes time-lapse images captured between 96 and 112 hpi, and the maternal age. Two separate models are incorporated to make predictions, one trained to discriminate between euploid (EUP) and aneuploid (ANU) embryos, and another trained to distinguish between euploid and complex aneuploid (CxA) embryos. The output from the model includes probabilities for euploidy, aneuploidy, and complex aneuploidy, along with intermediary quality scores that can be leveraged for further analysis of the embryo. The STORK-V platform serves as a valuable tool for embryologists and in vitro fertilization (IVF) clinics. It offers a convenient and efficient way to assess an embryo’s ploidy status, which is a crucial factor in the successful outcomes of assisted reproductive treatments. This will help medical professionals make more informed decisions regarding embryo selection and ultimately improve IVF success rates.

**Figure 5.**
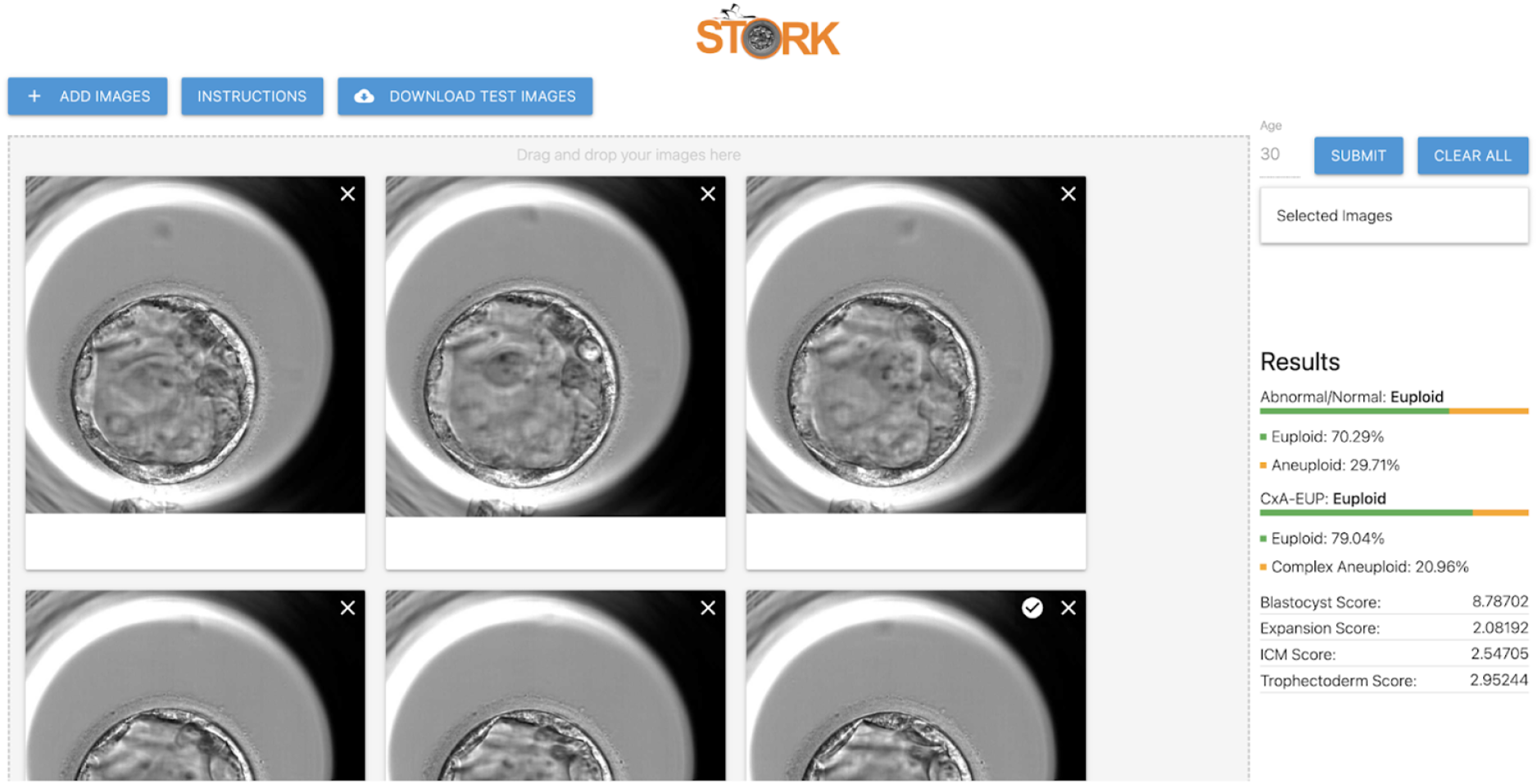
STORK-V web interface. A clinical tool that utilizes automation to assist embryologists in determining both the embryo quality score and ploidy status, providing a comprehensive assessment of the embryo.

## Discussion

In this study, we introduced BELA, which surpasses the traditional IVF embryo classification methods that usually rely on training data from later stages of embryo development and focus only on either image or video data. BELA stands out as a fully automated model that predicts blastocyst score and utilizes these predictions as a proxy for ploidy classification. BELA’s performance is competitive with a model trained on embryologist-annotated blastocyst scores and it significantly surpasses models trained exclusively on time-lapse imaging sequences without a proxy score. Remarkably, BELA only needs time-lapse images from 96 to 112 hpi and maternal age to predict an embryo’s ploidy status, thereby making it effortlessly adaptable to clinical workflows without causing any disruption. Notably, BELA also offers a degree of explainability; embryologists can use the model-derived blastocyst score (MDBS) and other scores predicted via multitasking to comprehend the rationale behind a specific ploidy status classification. In terms of recall, BELA demonstrates substantial potential for successfully selecting euploid embryos, especially for the WCM-Embryoscope+ dataset (**Supplemental Table 1**). While the model’s performance decreases in test datasets outside Weill Cornell, BELA still outperforms models trained on maternal age and/or embryologist-derived blastocyst score. BELA also interestingly found that single aneuploid embryos were evenly predicted as either euploid (EUP) or complex aneuploid (CxA) by our EUP versus CxA BELA model, suggesting that single aneuploid embryos often resemble euploid or complex aneuploid embryos, thus making their identification more challenging. These results are further confirmed by BELA models specifically trained to discriminate between euploid and single aneuploid embryos (**Supplemental Text 2). Supplemental Table 2** shows BELA’s AUC performance across various age groups classified by the Society for Assisted Reproductive Technology (SART). Despite maternal age being a strong predictor, performances across SART age groups tend to be bimodal (performing best at lower and higher age groups) for the WCM-Embryoscope and WCM-Embryoscope+ datasets. Moreover, in the Spain and Florida datasets, performances across maternal age do not follow the same distributions that were present in WCM datasets suggesting that these clinics’ varying demographics may affect model performance. In conclusion, while BELA is not intended to replace PGT-A, it can provide valuable supplementary information to support decision-making by embryologists, potentially leading to improved success rates in IVF procedures.

The study has several limitations. Firstly, video classification models, such as the one used in this study, demand substantial amounts of training data. Only ∼2,000 time-lapse sequences of embryo development were available for training, which restricted the ability to experiment with more computationally intensive video classification architectures like the 3D ConvNets or two-stream inflated convolutional nets. Secondly, despite trying multiple architectures for the feature extractor model, none performed as effectively as the ImageNet pretrained VGG16 architecture. There could potentially be more suitable feature extractors we did not consider, which might yield information from earlier stages of embryo development. Thirdly, we did not have access to several relevant maternal features, such as hormone levels at the time of oogenesis, demographics, and other clinically pertinent data. These could enhance the prediction of embryo ploidy status. Another limitation was the use of blastocyst scores as intermediary labels in BELA. Despite being well-documented, the blastocyst score is a manually curated label and can be subject to intra-observational bias. Nonetheless, we demonstrated that blastocyst score remains predictive of ploidy, justifying its use as an intermediary proxy value. The results might also be influenced by differing inclusion-exclusion criteria between datasets, possibly explaining some of the differences in model performance among the test datasets. After conducting a preliminary analysis (**Supplemental Text 3)**, we have developed the BELA model to not consider mosaic embryos and as such, mosaic embryos with high implantation potential could be misclassified. Regarding the ploidy status labels, the use of different platforms for PGT-A across clinics might impact the model’s accuracy and generalizability. There is significant variability in PGT-A results between labs and platforms, with no industry-wide standardization currently in place.^17^ Factors like methods used for biopsy preparation and the interpretation of results by clinicians could influence PGT-A results, possibly leading to differing detection rates of single versus complex aneuploidy.^18^ However, for the advancement of assistive reproductive technologies in IVF, the benchmark should be hastening the time to pregnancy and enhancing live birth outcomes. Embryo selection remains pivotal to this goal, necessitating the prioritization of embryos with high implantation potential and the de-prioritization of those with low potential. While most current embryo selection methodologies, such as morphological assessments, lack standardization and are largely subjective, PGT-A offers a consistent approach. This consistency is imperative for developing universally applicable embryo selection methods. Consequently, we used PGT-A results as our model’s ground truth labels. BELA aims to deliver a standardized, non-invasive, cost-effective, and efficient embryo selection and prioritization process. Lastly, the study’s model relies predominantly on data from time-lapse microscopy. Consequently, clinics lacking access to this technology will be unable to utilize the developed models.

Contrary to many prior studies that used non-viable embryos as negatives, leading to higher AUCs, the models developed in this study only consist of good biopsied embryos, making them more clinically applicable. The practical implications of these findings could significantly impact the efficiency and effectiveness of the embryo selection process. While the models developed, including BELA, do not replace PGT-A, they can help embryologists reduce the time and effort required to assess embryos. This streamlining of the workflow could allow faster decision-making, letting clinicians concentrate more on patient care and management. Using BELA could also decrease costs for patients and minimize risks associated with the biopsy process. By preselecting embryos with higher odds of being euploid, unnecessary biopsies can be avoided, reducing potential damage to the embryos and increasing the chances of a successful pregnancy. By deselecting embryos that are likely to be complex aneuploid, clinicians can gain insights into embryos that may cause embryo arrest or implantation failure. This is especially crucial for patients with a limited number of embryos, as it helps maximize the odds of success while minimizing potential risks. Furthermore, BELA can serve as a valuable tool for patient counseling and decision-making. With a clearer understanding of the embryo ploidy status, patients can make more informed decisions about their treatment options, considering the potential risks and benefits of each choice. Such increased transparency and patient involvement could potentially lead to improved patient satisfaction and trust in the IVF process. Automated blastocyst score prediction (MDBS) is also clinically relevant to embryologists currently manually annotating embryo scores. In situations where BELA is not used end-to-end to predict embryo ploidy, it could supplement manual embryo quality scoring. Additionally, BELA could standardize blastocyst scoring across clinics by providing an objective score free from embryologist subjectivity. Future iterations of models like BELA, which require no manually curated features and are fully automated from end-to-end, could be adopted into clinical practice.

## Materials & Methods

The research utilized multiple datasets for training and validation of the machine learning models. The first dataset, known as the WCM-Embryoscope data, was collected from the Center for Reproductive Medicine at Weill Cornell Medicine between 2018 and 2019. It comprises time-lapse images and PGT-A results for 1,998 embryos, including 494 single aneuploid (SA), 588 complex aneuploid (CxA), and 916 euploid (EUP) embryos. A total of 498 patients were included in the WCM-Embryoscope data with an average of four biopsied embryos each. We treated each sample independently irrespective of parental origin. Accompanying the time-lapse sequences were clinical data such as embryologist-derived blastocyst score (BS), morphokinetic parameters, and maternal age at the time of oocyte retrieval. The blastocyst score is the sum of a set of scores converted from the expansion, inner cell mass (ICM), trophectoderm (TE) grades, and day of blastocyst formation.^14^ The blastocyst score ranges from 3 to 14, with a lower number indicating a higher-quality embryo. The images were captured using the Embryoscope® imaging instrument. To validate the models’ generalizability, we used a second dataset, referred to as the WCM-Embryoscope+ data, which was also collected from the Center for Reproductive Medicine. However, these were gathered between 2019 and 2020 and included a total of 841 embryos (170 SA, 261 CxA, and 410 EUP), using a newer Embryoscope+® instrument. Similar to the first dataset, this also contained BS, morphokinetic parameters, and maternal age for each embryo. Furthermore, two external datasets were employed for further validation. The first, referred to as the Spain dataset, came from IVI Valencia and contained 543 embryos (309 ANU and 234 EUP) with time-lapse sequences, morphokinetic parameters, and maternal age. These images were also captured using the Embryoscope instrument. The second external dataset, referred to as the Florida dataset, was collected from IVF Florida and included 869 embryos (202 SA, 222 CxA, and 445 EUP) with maternal age and blastocyst score for each embryo. These images were captured using the Embryoscope+® instrument.

### Preimplantation Genetic Testing

Embryos from Weill Cornell were biopsied on Day 5 or Day 6 depending on when they reached blastocyst stage. Biopsied cells were analyzed using next generation sequencing (NGS) technology at the Ronald O. Perelman and Claudia Cohen Center for Reproductive Medicine (CRM). Analyses for the Spain Dataset were done by Igenomix Spain. Embryos were subjected to assisted hatching on Day 3, after cell counting, with the Hamilton-Thorne LykosVR laser. After reaching blastocyst stage, 5–6 trophectodermal cells were biopsied and their ploidy was assessed by Thermo Fisher Scientific’s NGS technology. Embryos from IVF Florida were also analyzed by Igenomix using Thermo Fisher Scientific’s NGS technology. More details about PGT-A protocols can be found in García-Pascual et al.^20^

### Temporal and Spatial Processing

Extracted time-lapse image sequences were highly variable in length, frame rate, start, and end points. These variabilities resulted in numerous embryos missing information from particular time periods, and a lack of proper annotation could lead to bias in model training. To mitigate these biases, the following protocol was developed to clean and standardize all time-lapse sequences, shown below.

1. Standardized timepoints are designated at 30-minute intervals from 0 to 150 hpi (i.e., 0 hpi, 0.5 hpi, … 149.5 hpi, 150.0 hpi).
2. For each embryo, time-lapse images taken closest to standardized time points are assigned to each standpoint. If there is no image close enough (within 2hrs) to the standardized time point, a blank frame is assigned to the standardized time point. We chose a 2-hour boundary as the ‘close enough’ range for several reasons. First, our observations indicated that significant changes in the embryos typically occurred at intervals greater than 2 hours. As a result, a 2-hour window provided a balance between accurately capturing significant changes while also allowing for reasonable data standardization. This timeframe was also influenced by the overall rate of data acquisition, which sometimes varied but was generally frequent enough to capture changes within this 2-hour window. However, we recognize the potential for variability and further studies may explore the impact of different time boundaries. At this point, each standardized time-lapse sequence has 301 frames, with each frame corresponding to a standardized test point between 0 and 150 hpi.
3. After construction of standardized time-lapse sequences, frames can be extracted for video classification model development using three parameters: start hr, end hr, and interval. For example, a model trained on day-2 embryo development would use these parameters, start hr = 24.0 hpi, end hr = 48.0 hpi, and interval = 2 hrs. This results in 13 frames.
4. For image classification tasks, a time point of focus can be ascertained and the frame assigned to that time point can be extracted.

We standardized the lengths, start, and end points of all time-lapse videos using set time points and intervals. Adjacent frames were utilized to impute missing time points. Some sequences, rendered unusable for certain prediction tasks post-standardization, were excluded from the analysis based on exclusion criteria. These criteria encompass instances where the embryo was absent from the petri dish, the embryo was less than half-visible, or the image was too dim to discern the embryo. We resized each frame from 800 × 800 to 224 × 224. To curtail background bias during model training, we implemented a circle Hough Transform for embryo segmentation in each video frame. This processing was uniformly applied across WCM-Embryoscope, WCM-Embryoscope+, Spain, and Florida datasets. To bolster the diversity and robustness of our training data, we incorporated video augmentation techniques, including random horizontal flipping and rotations. The former yielded mirror images of original frames, effectively doubling our data and fostering diverse pattern learning. Random rotations enhanced the model’s adaptability to varied embryo orientations, thereby simulating real-world scenarios. We opted for these techniques as they accurately represent potential real-world variations, fortifying our model’s robustness. Consequently, these augmentations improved our model’s generalization capability and enhanced its performance on unseen data.

### General Study Architecture

Two different prediction tasks were modeled between euploid (EUP), aneuploid (ANU), and complex aneuploid (CxA): EUP versus ANU and EUP versus CxA. Spatial features for each frame were extracted from the cleaned time-lapse images of the embryos using an ImageNet pretrained VGG16 convolutional neural network (CNN). Time-lapse image frames from 96 hpi to 112 hpi (day 5) were processed according to the Temporal and Spatial Processing section. The features extracted from these frames were input to a multi-task BiLSTM regression model (video regression task), which was primarily trained to predict embryologist-derived blastocyst scores. We investigated various dataset combinations for training the BELA models (**Supplemental Text 4**), ultimately using only WCM-Embryoscope data for the final models. To prevent data leakage, the WCM-Embryoscope dataset was split 70/30 for training/testing. This process exclusively utilized embryos which passed our exclusion criteria, reducing the dataset from 1,998 to 1,684 embryos. The BiLSTM regression model was trained only using the training slice of the dataset. Four-fold cross-validation was employed when training the BiLSTM regression models, setting aside data for monitoring validation loss. The predicted blastocyst scores for the training split embryos from the BiLSTM regression model, along with maternal age, were used to train a logistic regression model to predict embryo ploidy. A logistic regression model was trained on each of the cross-validated BiLSTM regression models, and the performance metrics of each logistic regression model were averaged. Model performance was measured using accuracy, area-under-receiver-operator-curve (AUC), precision, and recall.

We used the Student’s *t-test* to compare the means between two groups. This statistical test was selected because it is well-suited for comparing the means of two samples when the data is approximately normally distributed and the variances of the two groups are similar, as is the case with our data. In addition, all experiments were adjusted for multiple testing using Bonferroni correction to control for the increased chances of observing a statistically significant result due to the numerous tests performed.

### Feature Extraction

To extract spatial features from each frame of time-lapse images, an ImageNet pre-trained model from Tensorflow 2.7 was utilized. After experimenting with various pre-trained feature weights and extractors, we utilized a VGG16 CNN architecture to extract spatial features from images. The final layer of the pre-trained architecture performed average pooling which resulted in 512-dimensional feature vectors for each frame of each embryo.

### BELA Prediction Models

A BiLSTM network was employed for blastocyst score regression, leveraging its capabilities in sequential data pattern recognition, thus processing temporal information from time-lapse images.^19^ Our architecture comprises a bidirectional LSTM layer and three dense layers. The BiLSTM received 512-dimensional feature vectors extracted per frame for each embryo. While attention mechanisms and multiple bidirectional LSTM layers were explored, they failed to enhance performance significantly (p > 0.05) across all tasks. We modified the BiLSTM architecture to perform multitasking, wherein, in addition to the blastocyst score, the model was trained to predict expansion score, ICM score, and TE score. Multitasking has been used in previous studies to increase performance in scenarios where predicting different scenarios together may be advantageous to individual task performance. Similar tasks may have overlap in model weights required to come to accurate predictions, hence providing additional information for performing each task.^21^ Because expansion, ICM, and TE score make up the overall blastocyst score, we believe that multitasking can be used to improve blastocyst score prediction. The BiLSTM architecture consists of one bidirectional LSTM layer followed by two multi-unit dense layers. For each prediction task, a 1-unit dense layer is added to the model. Since all tasks of the multitask model are regression-based, we used”logcosh” as the loss function and Adam as the optimizer. Loss weights for each prediction task within the multitask environment were equal. Maternal age was included as a feature in the BiLSTM regression model to predict blastocyst score. Early-stopping with patience = 5 was used to ensure that the model was not overfitting to the training data by monitoring the validation loss on the cross-fold validation data. The performance of the first component of BELA was evaluated using the mean absolute error (MAE) of the predicted blastocyst score (MDBS). Multitask BELA demonstrated a lower MAE (1.855 ± 0.03) compared with a non-multitask BELA (1.877 ± 0.027) on the WCM-Embryoscope test, supporting the use of multitasking. The second part of BELA, the logistic regression model, was fed the predicted blastocyst score, sometimes in combination with maternal age, and performed a binary classification task. The logistic regression model used cross-entropy loss.

### Computational Resources and Time Requirements

Model training and inference were conducted using an Apple M1 Mac with TensorFlow Metal. Logistic regression models demonstrated an average training time of 2.5 ± 1.2 seconds, whereas BiLSTM models required 30.3 ± 11 minutes. The BELA model on the STORK-V platform was trained on a high-performance BioHPC computing cluster at Cornell, Ithaca, utilizing an NVIDIA A40 GPU and achieving a training time of 5.23 minutes. Inference for a single embryo on the STORK-V platform took 30 ± 5 seconds. The efficient use of consumer-grade hardware highlights the practicality of our models for assisted reproductive technology applications.

## Supporting information

All Supplemental Files

## Data Availability

The datasets analyzed in this study are not publicly available due to reasonable privacy and security concerns. The data were collected from multiple medical centers and have been shared with researchers only within the context of institutional review-board approved research collaborations. However, it is important to note that the method we developed is not specific to the datasets used in this study and can be applied to other imaging data sets. To that end, we have made our deep-learning model, BELA, available through a web-based user interface (https://stork-v.eipm-research.org/). Access to this password-protected site is granted for research purposes only and can be obtained by contacting the corresponding author.

## Code Availability

Code used to train and evaluate the models can be found at https://github.com/eipm/stork-v/.

## Acknowledgements

I.H is supported by an NIGMS Maximizing Investigators’ Research Award (MIRA) R35GM138152. The content is solely the responsibility of the authors and does not necessarily represent the official views of the National Institutes of Health. Through allocation TG-ASC190055 to I.H., this work used the Extreme Science and Engineering Discovery Environment (XSEDE), which is supported by National Science Foundation grant number ACI-1548562. SR would like to acknowledge the support from Tri-Institutional Training Program in Computational Biology and Medicine (CBM) funded by the NIH grant 1T32GM083937.

## Author Contributions

S.R., J.B., N.Z., and I.H. conceived the study. S.R., J.B., M.B., and I.H. conceived the method and designed the algorithmic techniques. S.R. wrote the codes and performed the computational analysis with input from I.H, J.B., M.B., and K.O.

Q.Z., J.E.M., Z.R., and N.Z. provided the Weill Cornell datasets and labeled images. N.Z. evaluated additional embryo images. M.M. provided the Spain dataset. K.A.M. and D.H. provided the Florida dataset. S.R. drafted the manuscript with input from J.B., M.B., O.E., Q.Z., and I.H. P.Z. and A.S. designed the user interface. All the authors read the paper and suggested edits. I.H. supervised the project.

