## Supplementary material for "Automatic Ploidy Prediction and Quality Assessment of Human Blastocyst Using Time-Lapse Imaging": All Supplemental Files

### Supplemental Information

#### S1. Supplemental Tables

| Test Set | Prediction Task | Prediction Setting | AUC | Recall | Precision |
|---|---|---|---|---|---|
| WCM-Embryoscope | EUP vs. CxA | BELA | 0.708 ± 0.004 | 0.591 ± 0.019 | 0.797 ± 0.019 |
| | | BELA + Age | 0.826 ± 0.004 | 0.745 ± 0.016 | 0.878 ± 0.008 |
| | EUP vs. ANU | BELA | 0.659 ± 0.008 | 0.546 ± 0.009 | 0.643 ± 0.027 |
| | | BELA + Age | 0.76 ± 0.002 | 0.688 ± 0.01 | 0.705 ± 0.005 |
| WCM-Embryoscope+ | EUP vs. CxA | BELA | 0.67 ± 0.001 | 0.842 ± 0.013 | 0.668 ± 0.001 |
| | | BELA + Age | 0.771 ± 0.002 | 0.879 ± 0.004 | 0.722 ± 0.006 |
| | EUP vs. ANU | BELA | 0.626 ± 0.001 | 0.813 ± 0.027 | 0.536 ± 0.001 |
| | | BELA + Age | 0.717 ± 0.002 | 0.85 ± 0.003 | 0.594 ± 0.004 |
| Spain | EUP vs. ANU | BELA | 0.648 ± 0.001 | 0.679 ± 0.014 | 0.524 ± 0.012 |
| | | BELA + Age | 0.711 ± 0.001 | 0.699 ± 0.016 | 0.571 ± 0.005 |
| Florida | EUP vs. CxA | BELA | 0.591 ± 0.001 | 0.625 ± 0.021 | 0.709 ± 0.003 |
| | | BELA + Age | 0.651 ± 0.001 | 0.787 ± 0.015 | 0.719 ± 0.001 |
| | EUP vs. ANU | BELA | 0.572 ± 0.001 | 0.583 ± 0.024 | 0.556 ± 0.003 |
| | | BELA + Age | 0.621 ± 0.002 | 0.751 ± 0.007 | 0.570 ± 0.004 |

**Supplemental Table 1. All performance metrics for BELA on all prediction tasks, settings, and test sets.** AUC, precision, and recall are shown. Within the prediction settings, we show performances of a BELA model that only uses MDBS and a BELA model that uses both maternal age and MDBS.

| SART Age Group | WCM-Embryoscope AUC | WCM-Embryoscope + AUC | Spain AUC | Florida AUC |
|---|---|---|---|---|
| < 35 | 0.686 ± 0.019 (112) | 0.590 ± 0.001 (188) | 0.685 ± 0.002 (105) | 0.541 ± 0.003 (321) |
| 35 - 37 | 0.650 ± 0.008 (138) | 0.604 ± 0.001 (272) | 0.607 ± 0.001 (123) | 0.504 ± 0.002 (234) |
| 38 - 40 | 0.534 ± 0.006 (120) | 0.580 ± 0.001 (226) | 0.619 ± 0.001 (182) | 0.632 ± 0.001 (232) |
| 41 - 42 | 0.635 ± 0.044 (67) | 0.677 ± 0.001 (100) | 0.602 ± 0.001 (114) | 0.529 ± 0.002 (56) |
| > 42* | 0.548 ± 0.022 (59) | 0.593 ± 0.002 (55) | 0.912 ± 0.001 (19) | 0.864 ± 0.001 (26) |

**Supplemental Table 2. Performance of BELA on discriminating between euploid and aneuploid embryos across SART age groups.** The table shows the AUC performance of the BELA on the EUP versus ANU task across all tests and SART age groups is shown. The sample size in each age group for that dataset is shown in parentheses. *Group age > 42 has a significantly smaller sample size compared with other age groups which may increase the sensitivity of results within that age group.

| Timepoint | Video Classification Parameters | Image Classification Parameters | Embryoscope Data Distribution | Embryoscope+ Data Distribution |
|---|---|---|---|---|
| **Day 1** | *start*=6 hpi, *end*=24 hpi, *interval*=2 hrs | *extracted timepoint*=12 hpi | SA: 414, CxA: 456, EUP: 829 | SA: 169, CxA: 252, EUP: 402 |
| **Day 2** | *start*=24 hpi, *end*=48 hpi, *interval*=2 hrs | *extracted timepoint*=36 hpi | SA: 425, CxA: 469, EUP: 848 | SA: 168, CxA: 257, EUP: 408 |
| **Day 3** | *start*=48 hpi, *end*=72 hpi, *interval*=2 hrs | *extracted timepoint*=60 hpi | SA: 425, CxA: 467, EUP: 847 | SA: 169, CxA: 261, EUP: 410 |
| **Day 4** | *start*=72 hpi, *end*=96 hpi, *interval*=2 hrs | *extracted timepoint*=84 hpi | SA: 425, CxA: 465, EUP: 846 | SA: 170, CxA: 261, EUP: 410 |
| **day-5** | *start*=96 hpi, *end*=112 hpi, *interval*=2 hrs | *extracted timepoint*=108 hpi | SA: 415, CxA: 457, EUP: 812 | SA: 170, CxA: 261, EUP: 410 |

**Supplemental Table 3. Criteria and parameters for different prediction settings across time periods.** Start, end, and interval parameters are provided for video classification tasks. Time points at which frames were extracted for image classification are shown. Day 1 and day 5 have relatively shorter parameter ranges (18hrs, 16hrs) to account for missing data at extremely early and late stages of development. Data distributions for WCM-Embryoscope and WCM-Embryoscope+ data at different time periods are also shown. For WCM-Embryoscope data, distributions shown are before data augmentation.

| Discrimination Task | Prediction Setting | Metric | Day 1 | Day 2 | Day 3 | Day 4 | Day 5 |
|---|---|---|---|---|---|---|---|
| EUP vs. ANU | Video | AUC | 0.514 ± 0.016 | 0.538 ± 0.007 | 0.522 ± 0.01 | 0.499 ± 0.006 | 0.564 ± 0.013 |
| | | Precision | 0.546 ± 0.027 | 0.534 ± 0.013 | 0.542 ± 0.003 | 0.491 ± 0.049 | 0.585 ± 0.028 |
| | | Recall | 0.473 ± 0.303 | 0.767 ± 0.288 | 0.37 ± 0.152 | 0.459 ± 0.247 | 0.403 ± 0.106 |
| | Image | AUC | 0.513 ± 0.01 | 0.518 ± 0.014 | 0.515 ± 0.004 | 0.507 ± 0.004 | 0.545 ± 0.008 |
| | | Precision | 0.527 ± 0.011 | 0.535 ± 0.012 | 0.532 ± 0.005 | 0.524 ± 0.008 | 0.547 ± 0.007 |
| | | Recall | 0.482 ± 0.012 | 0.475 ± 0.008 | 0.479 ± 0.021 | 0.459 ± 0.018 | 0.483 ± 0.011 |
| | Video + Age | AUC | 0.719 ± 0.004 | 0.732 ± 0.004 | 0.724 ± 0.009 | 0.726 ± 0.009 | 0.731 ± 0.006 |
| | | Precision | 0.673 ± 0.008 | 0.696 ± 0.008 | 0.676 ± 0.012 | 0.669 ± 0.021 | 0.699 ± 0.013 |
| | | Recall | 0.722 ± 0.039 | 0.675 ± 0.031 | 0.692 ± 0.027 | 0.717 ± 0.047 | 0.631 ± 0.032 |
| | Image + Age | AUC | 0.682 ± 0.014 | 0.702 ± 0.004 | 0.689 ± 0.006 | 0.688 ± 0.011 | 0.705 ± 0.004 |
| | | Precision | 0.657 ± 0.014 | 0.669 ± 0.012 | 0.651 ± 0.005 | 0.664 ± 0.01 | 0.677 ± 0.003 |
| | | Recall | 0.624 ± 0.01 | 0.655 ± 0.009 | 0.643 ± 0.01 | 0.639 ± 0.008 | 0.639 ± 0.005 |
| EUP vs. CxA | Video | AUC | 0.515 ± 0.004 | 0.523 ± 0.018 | 0.501 ± 0.018 | 0.505 ± 0.019 | 0.622 ± 0.014 |
| | | Precision | 0.689 ± 0.007 | 0.687 ± 0.012 | 0.673 ± 0.006 | 0.641 ± 0.078 | 0.805 ± 0.043 |
| | | Recall | 0.518 ± 0.212 | 0.65 ± 0.159 | 0.556 ± 0.175 | 0.496 ± 0.323 | 0.41 ± 0.149 |
| | Image | AUC | 0.508 ± 0.013 | 0.515 ± 0.009 | 0.529 ± 0.017 | 0.511 ± 0.013 | 0.571 ± 0.007 |
| | | Precision | 0.684 ± 0.006 | 0.69 ± 0.002 | 0.691 ± 0.003 | 0.692 ± 0.004 | 0.696 ± 0.002 |
| | | Recall | 0.845 ± 0.009 | 0.832 ± 0.008 | 0.851 ± 0.007 | 0.835 ± 0.001 | 0.807 ± 0.008 |
| | Video + Age | AUC | 0.772 ± 0.008 | 0.781 ± 0.008 | 0.778 ± 0.013 | 0.782 ± 0.008 | 0.795 ± 0.006 |
| | | Precision | 0.836 ± 0.017 | 0.849 ± 0.008 | 0.855 ± 0.011 | 0.852 ± 0.016 | 0.854 ± 0.015 |
| | | Recall | 0.739 ± 0.029 | 0.718 ± 0.032 | 0.712 ± 0.022 | 0.711 ± 0.037 | 0.71 ± 0.061 |
| | Image + Age | AUC | 0.775 ± 0.002 | 0.793 ± 0.007 | 0.764 ± 0.003 | 0.777 ± 0.009 | 0.784 ± 0.006 |
| | | Precision | 0.802 ± 0.008 | 0.809 ± 0.005 | 0.797 ± 0.007 | 0.811 ± 0.005 | 0.808 ± 0.005 |
| | | Recall | 0.843 ± 0.01 | 0.842 ± 0.008 | 0.827 ± 0.01 | 0.843 ± 0.009 | 0.834 ± 0.006 |

(Columns Day 1–Day 5 are under the spanning header "Time Point".)

**Supplemental Table 4. All performance metrics for prediction tasks on WCM-Embryoscope data.** AUC, precision, and recall on WCM-Embryoscope testing data for all prediction settings. Precision and recall were determined using a 0.5 threshold on predicted probability outputs.

| Discrimination Task | Prediction Setting | Metric | Time Point: Day 1 | Day 2 | Day 3 | Day 4 | Day 5 |
|---|---|---|---|---|---|---|---|
| EUP vs. ANU | Video | AUC | 0.489 ± 0.01 | 0.522 ± 0.018 | 0.487 ± 0.01 | 0.499 ± 0.02 | 0.577 ± 0.011 |
| | | Precision | 0.356 ± 0.206 | 0.503 ± 0.014 | 0.101 ± 0.176 | 0.355 ± 0.205 | 0.519 ± 0.017 |
| | | Recall | 0.379 ± 0.369 | 0.83 ± 0.213 | 0.018 ± 0.032 | 0.645 ± 0.399 | 0.709 ± 0.076 |
| | Image | AUC | 0.48 ± 0.006 | 0.473 ± 0.02 | 0.476 ± 0.002 | 0.503 ± 0.012 | 0.532 ± 0.011 |
| | | Precision | 0.471 ± 0.011 | 0.392 ± 0.07 | 0.432 ± 0.011 | 0.509 ± 0.014 | 0.5 ± 0.006 |
| | | Recall | 0.504 ± 0.038 | 0.074 ± 0.022 | 0.175 ± 0.036 | 0.26 ± 0.033 | 0.774 ± 0.015 |
| | Video + Age | AUC | 0.695 ± 0.008 | 0.704 ± 0.003 | 0.696 ± 0.006 | 0.698 ± 0.007 | 0.684 ± 0.014 |
| | | Precision | 0.639 ± 0.016 | 0.634 ± 0.005 | 0.639 ± 0.013 | 0.607 ± 0.013 | 0.596 ± 0.016 |
| | | Recall | 0.707 ± 0.055 | 0.722 ± 0.013 | 0.663 ± 0.048 | 0.762 ± 0.054 | 0.75 ± 0.03 |
| | Image + Age | AUC | 0.669 ± 0.007 | 0.658 ± 0.01 | 0.664 ± 0.012 | 0.665 ± 0.012 | 0.667 ± 0.012 |
| | | Precision | 0.61 ± 0.011 | 0.625 ± 0.018 | 0.632 ± 0.006 | 0.629 ± 0.014 | 0.572 ± 0.003 |
| | | Recall | 0.702 ± 0.025 | 0.388 ± 0.076 | 0.53 ± 0.036 | 0.491 ± 0.067 | 0.821 ± 0.015 |
| EUP vs. CxA | Video | AUC | 0.508 ± 0.015 | 0.505 ± 0.008 | 0.479 ± 0.067 | 0.487 ± 0.019 | 0.613 ± 0.004 |
| | | Precision | 0.712 ± 0.167 | 0.618 ± 0.004 | 0.572 ± 0.094 | 0.616 ± 0.011 | 0.658 ± 0.015 |
| | | Recall | 0.365 ± 0.323 | 0.92 ± 0.067 | 0.209 ± 0.176 | 0.53 ± 0.44 | 0.725 ± 0.041 |
| | Image | AUC | 0.503 ± 0.026 | 0.491 ± 0.017 | 0.481 ± 0.01 | 0.51 ± 0.01 | 0.568 ± 0.017 |
| | | Precision | 0.612 ± 0.007 | 0.591 ± 0.013 | 0.595 ± 0.015 | 0.614 ± 0.01 | 0.613 ± 0.004 |
| | | Recall | 0.89 ± 0.029 | 0.241 ± 0.014 | 0.421 ± 0.025 | 0.719 ± 0.126 | 0.958 ± 0.012 |
| | Video + Age | AUC | 0.747 ± 0.015 | 0.742 ± 0.006 | 0.732 ± 0.013 | 0.735 ± 0.016 | 0.741 ± 0.006 |
| | | Precision | 0.758 ± 0.015 | 0.767 ± 0.008 | 0.77 ± 0.012 | 0.745 ± 0.017 | 0.723 ± 0.02 |
| | | Recall | 0.785 ± 0.037 | 0.753 ± 0.013 | 0.662 ± 0.113 | 0.78 ± 0.028 | 0.826 ± 0.068 |
| | Image + Age | AUC | 0.738 ± 0.004 | 0.739 ± 0.006 | 0.721 ± 0.007 | 0.73 ± 0.008 | 0.732 ± 0.009 |
| | | Precision | 0.707 ± 0.006 | 0.772 ± 0.008 | 0.769 ± 0.011 | 0.733 ± 0.02 | 0.676 ± 0.017 |
| | | Recall | 0.86 ± 0.022 | 0.758 ± 0.009 | 0.726 ± 0.015 | 0.811 ± 0.032 | 0.926 ± 0.022 |

**Supplemental Table 5. All performance metrics for prediction tasks on WCM-Embryoscope+ data.** Shown above are AUC, precision, and recall on WCM-Embryoscope+ data for all prediction settings.

| Prediction Task and Dataset | day-5 AUC | All Days AUC |
|---|---|---|
| **EUP vs CxA, WCM-Embryoscope** | 0.622 ± 0.014 | 0.606 ± 0.011 |
| **EUP vs ANU, WCM-Embryoscope** | 0.564 ± 0.013 | 0.538 ± 0.025 |
| **EUP vs CxA, WCM-Embryoscope+** | 0.613 ± 0.004 | 0.589 ± 0.016 |
| **EUP vs ANU, WCM-Embryoscope+** | 0.577 ± 0.011 | 0.522 ± 0.041 |

**Supplemental Table 6. Video-only classification model performance between day 5 and all days.** AUC scores for all video-only prediction tasks on both WCM-Embryoscope and WCM-Embryoscope+ datasets. All-day models used the following parameters: *start*=6 hpi, *end*=112 hpi, and *interval*=2 hrs. Day-5 models perform significantly better than all-day models on all tasks and datasets.

| | BiLSTM AUC | XGBoost AUC |
|---|---|---|
| **EUP vs CxA, Video** | 0.622 ± 0.014 | 0.58 ± 0.02 |
| **EUP vs ANU, Video** | 0.564 ± 0.013 | 0.546 ± 0.009 |

**Supplemental Table 7. Day-5 video classification model Architecture comparisons.** AUC performances on the WCM-Embryoscope dataset with BiLSTM and XGBoost video classification day-5 models. Frame features for each embryo were concatenated before being inputted into XGBoost models. BiLSTM models perform significantly better than XGBoost models.

| Clinical Feature | WCM-Embryoscope | WCM-Embryoscope + | Spain | Florida |
|---|---|---|---|---|
| **Maternal Age** | -0.346 | -0.357 | -0.327 | -0.205 |
| **Blastocyst Score** | -0.306 | -0.254 | --- | -0.101 |

**Supplemental Table 8. Pearson correlations between clinical features and ploidy status.** Ploidy status was divided into Euploid and Aneuploid for this analysis. The Pearson correlation between clinical features and ploidy across all test sets is shown. No correlation between the blastocyst score and ploidy status is available for the Spain data because embryologist-derived blastocyst scores were not provided in the dataset.

| Prediction Task and Setting | 108 hpi AUC | 112 hpi AUC |
|---|---|---|
| **EUP vs CxA, Image** | 0.571 ± 0.007 | 0.57 ± 0.016 |
| **EUP vs ANU, Image** | 0.545 ± 0.008 | 0.552 ± 0.014 |
| **EUP vs CxA, Image + Age** | 0.784 ± 0.006 | 0.782 ± 0.009 |
| **EUP vs ANU, Image + Age** | 0.705 ± 0.004 | 0.706 ± 0.007 |

**Supplemental Table 9. Performance comparison of two day-5 image classification models.** AUC performances on the WCM-Embryoscope test set for day-5 image classification models. The model whose performance is shown in the middle column uses single images from 108 hpi and the right column uses single images from 112 hpi. Differences in performance are non-significant for all prediction tasks and settings.

| SART Age Group | WCM-Embryoscope | | WCM-Embryoscope + | | Spain | Florida | |
|---|---|---|---|---|---|---|---|
| **Categories** | Ploidy | BS | Ploidy | BS | Ploidy | Ploidy | BS |
| **< 35** | EUP: 262<br>SA: 74<br>CxA: 29 | 7.45 ± 2.52 | EUP: 131<br>SA: 30<br>CxA: 27 | 6.07 ± 1.94 | EUP: 67<br>ANU: 38 | EUP: 199<br>SA: 65<br>CxA: 57 | 7.53 ± 1.12 |
| **35 - 37** | EUP: 286<br>SA: 109<br>CxA: 52 | 7.8 ± 2.66 | EUP: 164<br>SA: 57<br>CxA: 51 | 6.02 ± 1.89 | EUP: 69<br>ANU: 54 | EUP: 118<br>SA: 58<br>CxA: 58 | 7.48 ± 1.19 |
| **38 - 40** | EUP: 159<br>SA: 110<br>CxA: 148 | 8.42 ± 2.67 | EUP: 83<br>SA: 54<br>CxA: 89 | 6.46 ± 1.93 | EUP: 70<br>ANU: 112 | EUP: 108<br>SA: 65<br>CxA: 59 | 7.42 ± 1.16 |
| **41 - 42** | EUP: 64<br>SA: 76<br>CxA: 111 | 8.37 ± 2.76 | EUP: 25<br>SA: 21<br>CxA: 54 | 6.39 ± 1.79 | EUP: 26<br>ANU: 88 | EUP: 16<br>SA: 10<br>CxA: 30 | 7.38 ± 1.0 |
| **> 42*** | EUP: 41<br>SA: 46<br>CxA: 117 | 9.02 ± 2.75 | EUP: 7<br>SA: 8<br>CxA: 40 | 6.87 ± 2.01 | EUP: 2<br>ANU: 17 | EUP: 4<br>SA: 4<br>CxA: 18 | 8.77 ± 1.95 |

**Supplemental Table 10. Distributions of ploidy and blastocyst score across SART age groups.** The table shows the distribution of embryo ploidy status and blastocysts score for all datasets across SART age groups. Spain embryos do not have SA/CxA discrimination and are all marked as ANU if they are aneuploid.

| Prediction Model | WCM-Embryoscope | WCM-Embryoscope+ | Florida |
|---|---|---|---|
| **Embryologist BS** | 0.579 ± 0.003 | 0.573 ± 0.004 | 0.506 ± 0.002 |
| **BELA** | 0.597 ± 0.008 | 0.575 ± 0.002 | 0.571 ± 0.003 |
| **Embryologist BS + Age** | 0.624 ± 0.003 | 0.644 ± 0.001 | 0.584 ± 0.003 |
| **BELA + Age** | 0.682 ± 0.005 | 0.648 ± 0.001 | 0.608 ± 0.004 |

**Supplemental Table 11. Performances of EUP vs SA models across tests.** AUC performances of EUP vs SA models (Embryologist BS and BELA) on all applicable test sets.

#### S2. Supplemental Figures

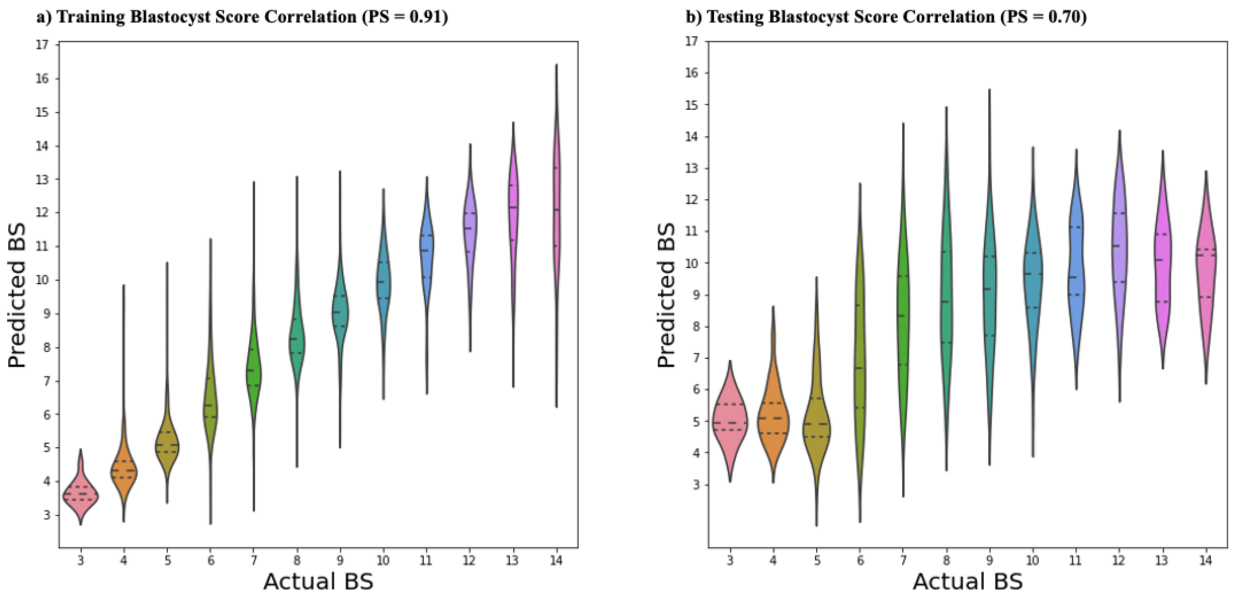


**Supplemental Figure 1. Correlations between predicted and actual BS scores from BELA.** Violin plots of score distributions of BELA model. Pearson correlation scores are shown in the panel titles (PS=*). (a) shows correlations within the training set. (b) Correlations within the WCM-Embryoscope test set.

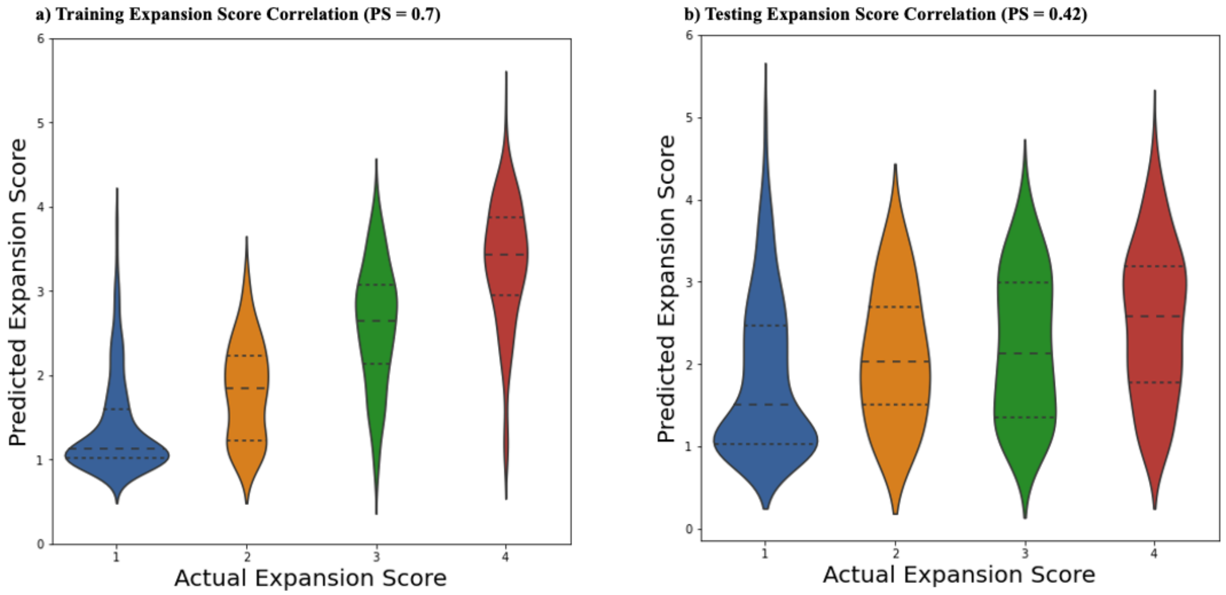

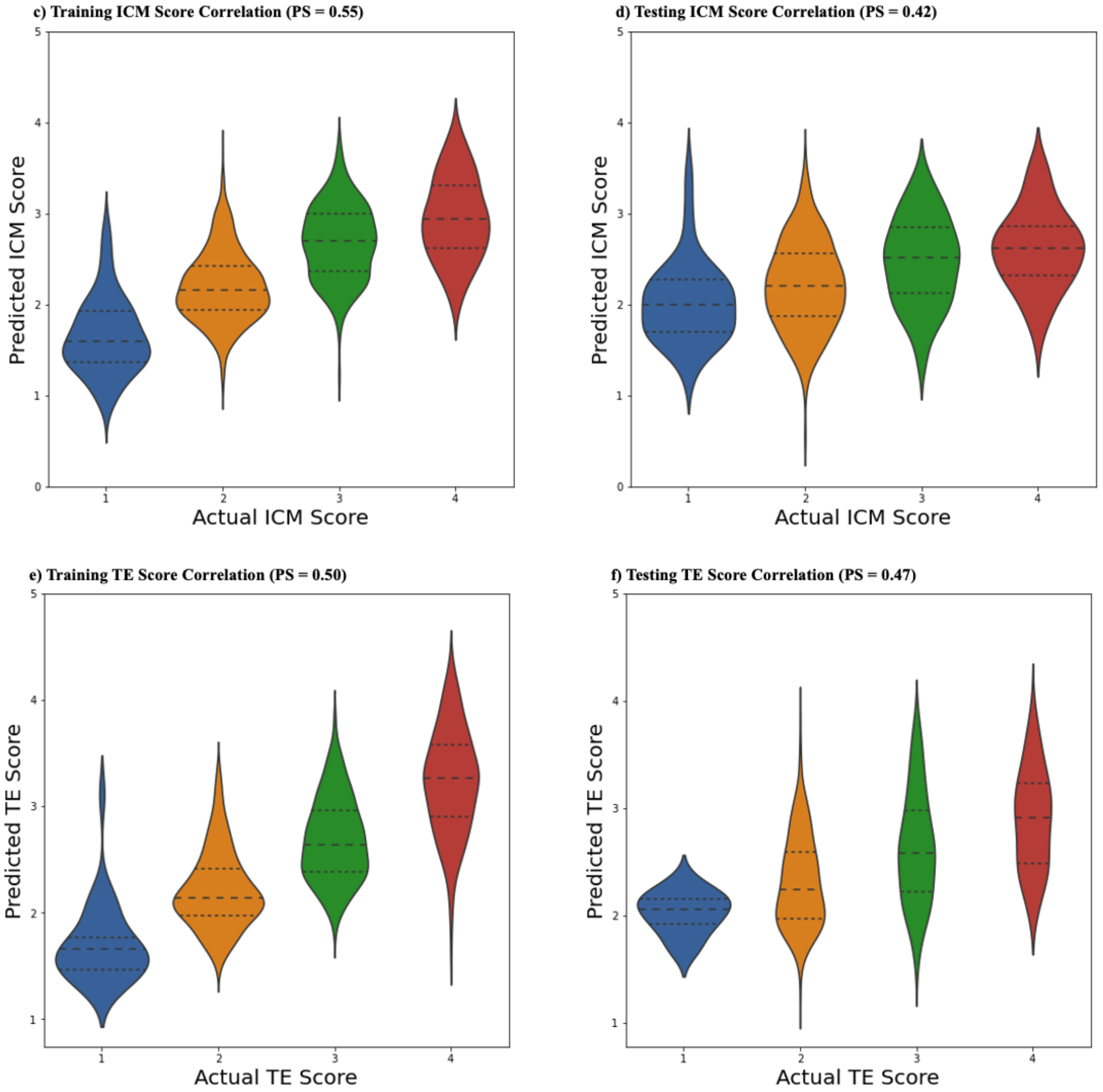


**Supplemental Figure 2. Correlations between predicted and actual multitask scores from BELA.** Violin plots of score distributions of BELA model. Correlation plots for other predicted scores in the first component of BELA (not used in downstream processes) are shown. Pearson correlations are shown in the panel titles (PS=*). (a, c, and e) Correlations within the training set. (b, d, and f) Correlations within the WCM-Embryoscope test set.

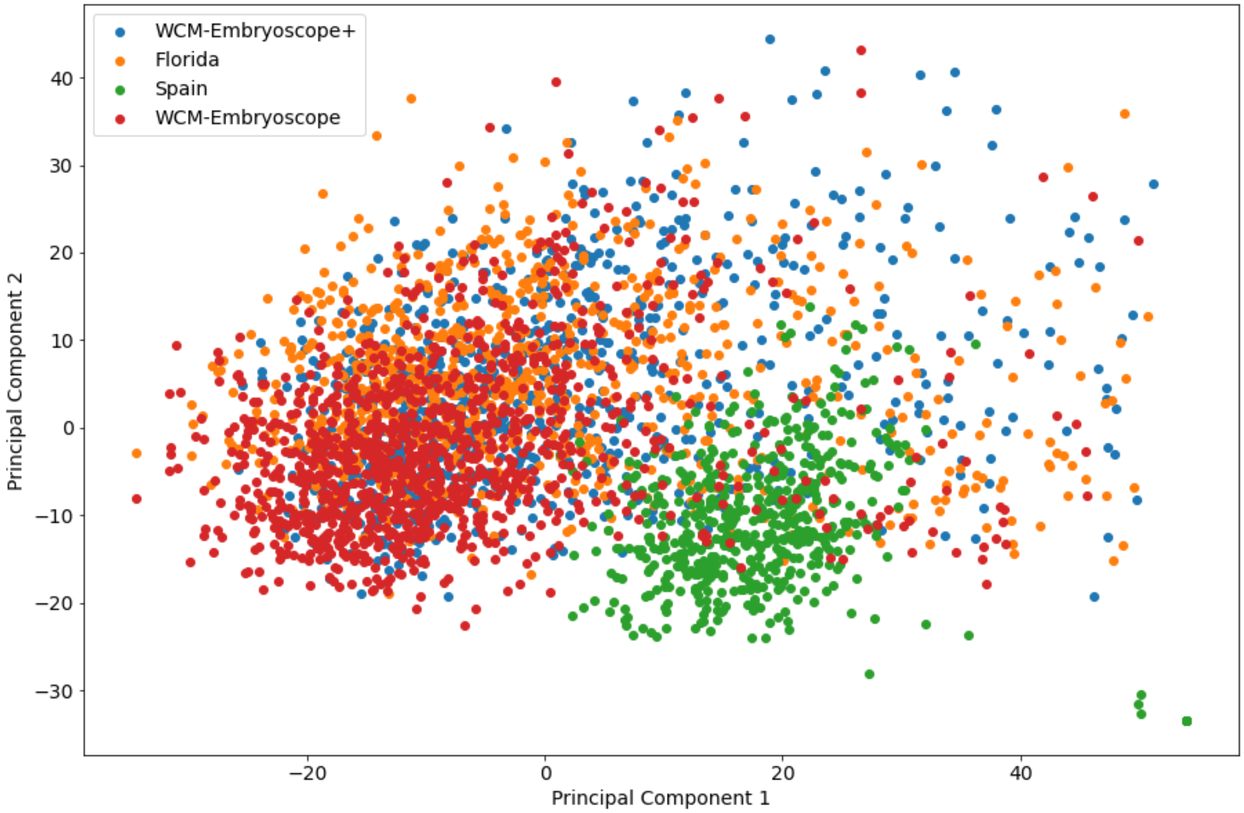


**Supplemental Figure 3. Feature encodings of embryo time-lapse videos by dataset.** Feature encodings were generated by applying PCA on feature vectors generated from a pretrained CNN model. Green points are part of the IVI Valencia Spain dataset. Orange points are part of the Florida dataset. Red points are part of the Weill Cornell Medicine internal dataset. Blue points are part of the WCM-Embryoscope+ dataset.

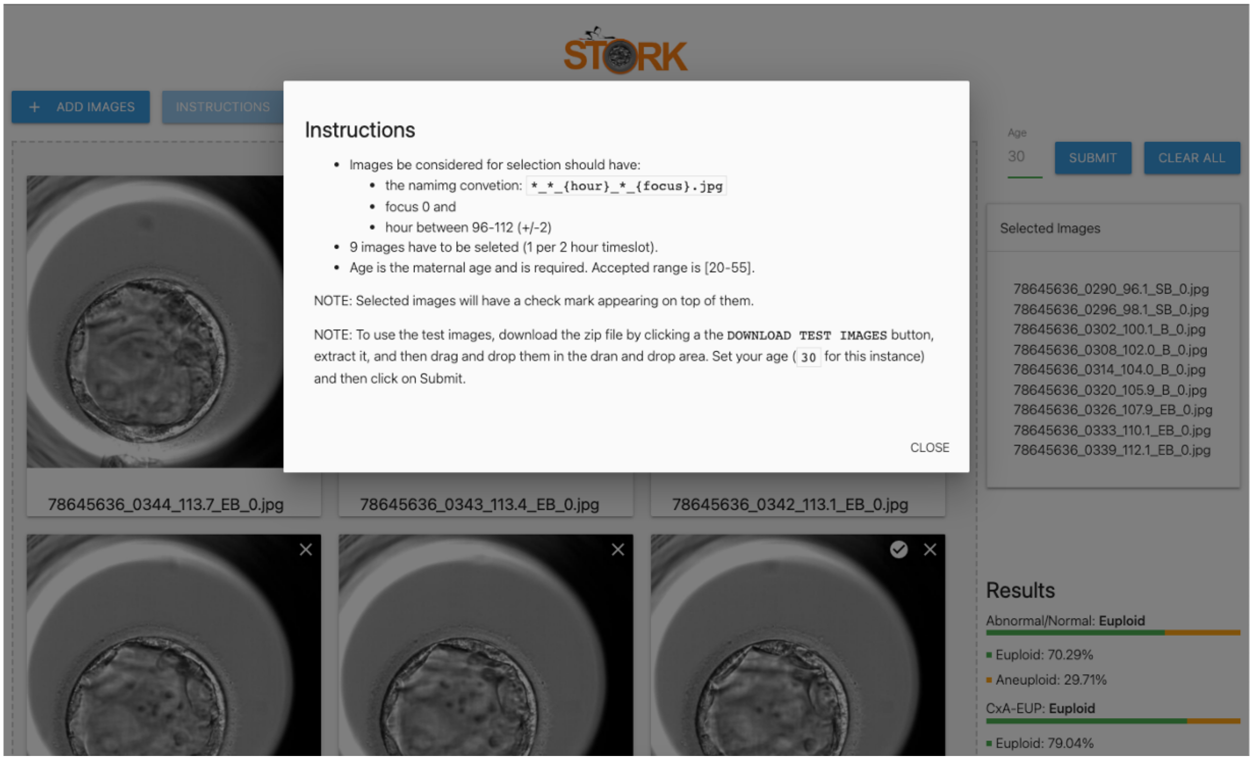


**Supplemental Figure 4. Instructions for using STORK-V.** Instructions for users to format time-lapse images for processing by STORK-V and analyzing test data.

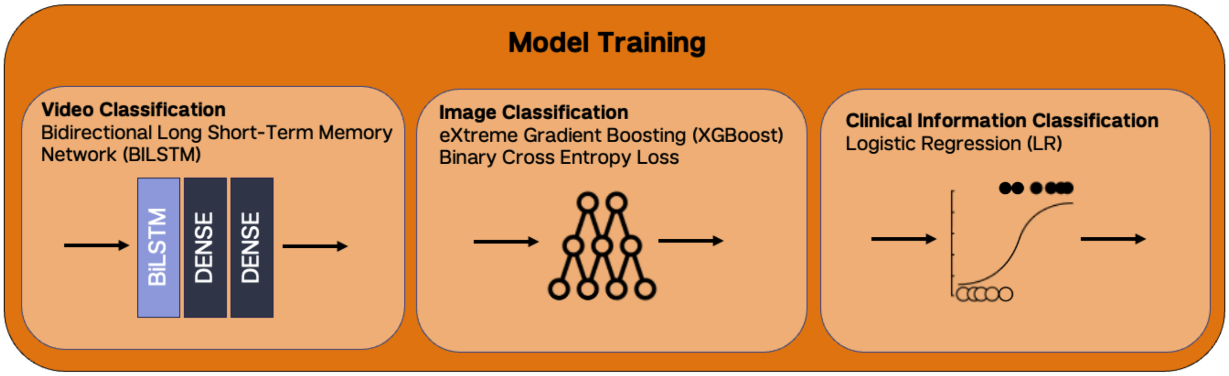


**Supplemental Figure 5. Overview of comparison analyses methodology.** Data is collected from WCM-Embryoscope and WCM-Embryoscope+ data sources. Preprocessing steps to create feature extractions are done via Figure 1 steps 1 - 4. A machine learning model is trained (only on WCM-Embryoscope data) to process the frame(s), where the architecture is dependent on whether the task is one of image or video classification.

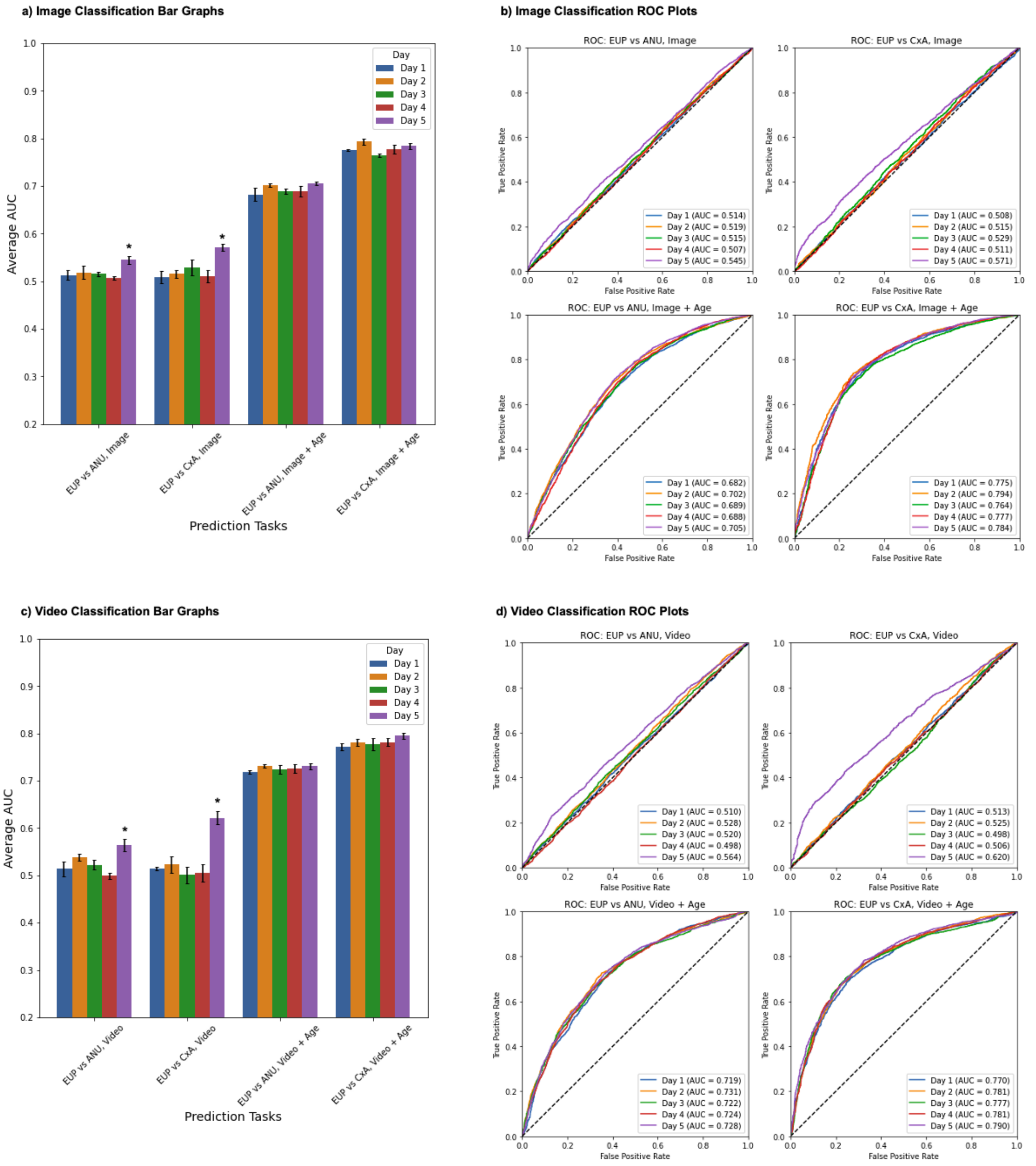


**Supplemental Figure 6**. **Model performances of classification models across time periods and prediction tasks.** (a) Average AUC with standard error bars across different days and different image classification tasks. (b) ROC plots for each image classification prediction task. AUC in ROC plots were calculated by stacking predictions across cross-folds for each group and then calculating an overall score. Each plot illustrates the performance of the prediction model in the given settings. (c) Average AUC with standard error bars across different days and different video classification tasks. (d) ROC plots for each video classification prediction task.

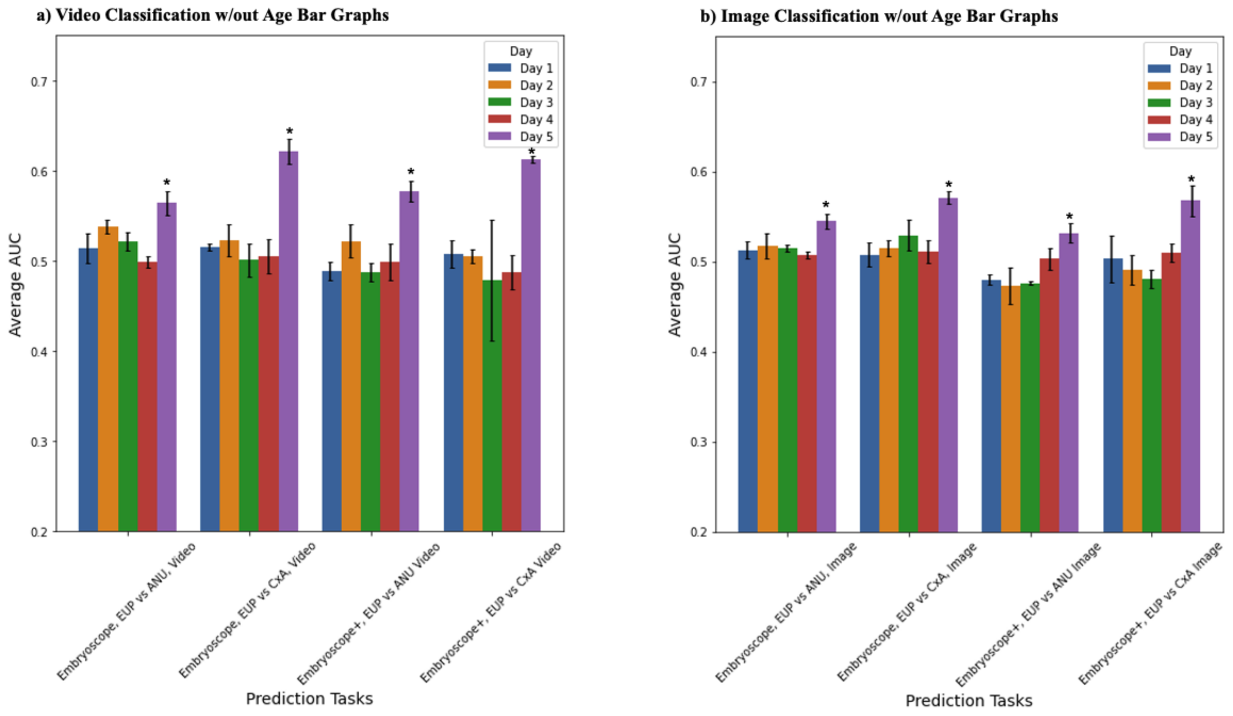


**Supplemental Figure 7**. **Model performances of classification models without maternal age across time periods on WCM-Embryoscope and WCM-Embryoscope+ datasets.** (a) Average AUC with standard error bars across different days and different video classification tasks. (b) Average AUC with standard error bars across different days and different image classification tasks.

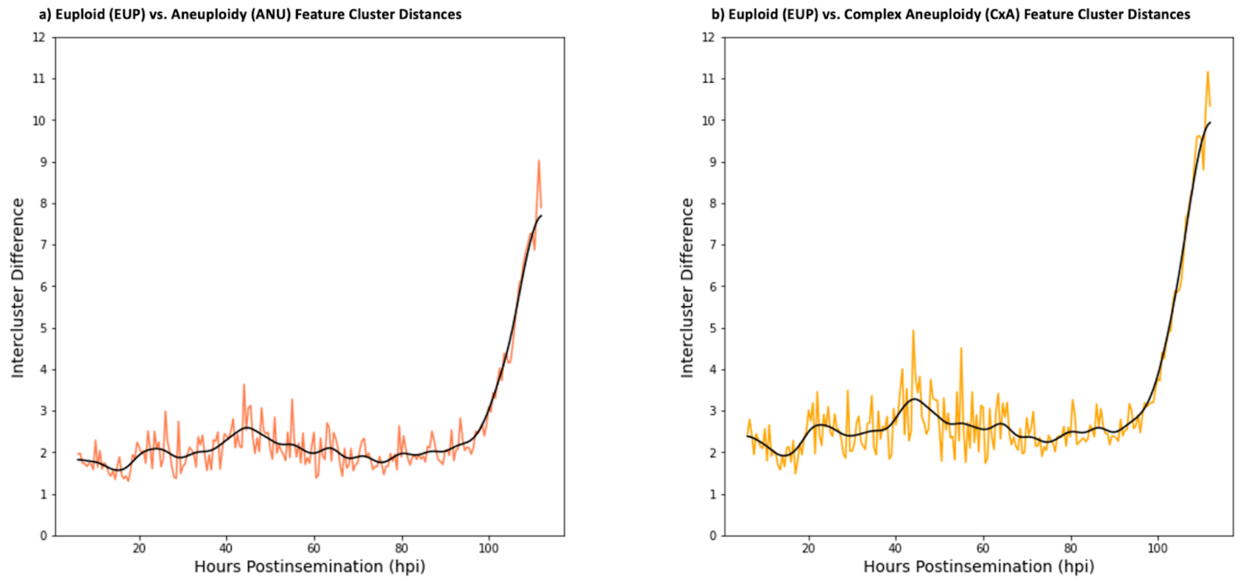

**Supplemental Figure 8**. **Intercluster differences between embryos clustered by ploidy status.** Intercluster differences, measured as the distance between cluster centroids where each cluster is a ploidy status, is shown for all WCM-Embryoscope embryos. Intercluster difference at each timepoint (i.e. hours post-insemination) is shown for both (a) EUP versus ANU and (b) EUP versus. CxA. Black lines are Gaussian smoothed curves that show the general trend of intercluster differences.

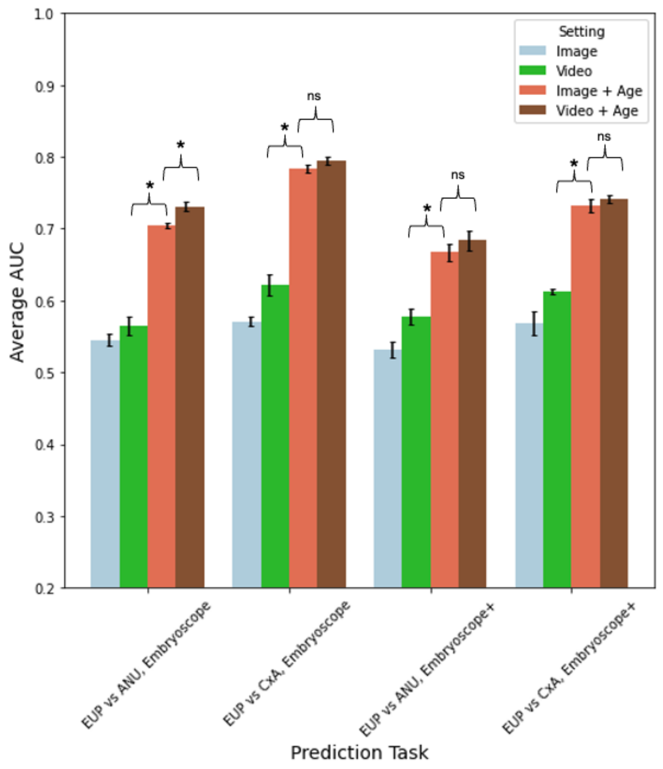


**Supplemental Figure 9**. **Model performances of day-5 image and video classification models.** Performances for both WCM-Embryoscope and WCM-Embryoscope+ datasets for all prediction tasks are depicted. Average AUC and standard errors are shown for each prediction setting.

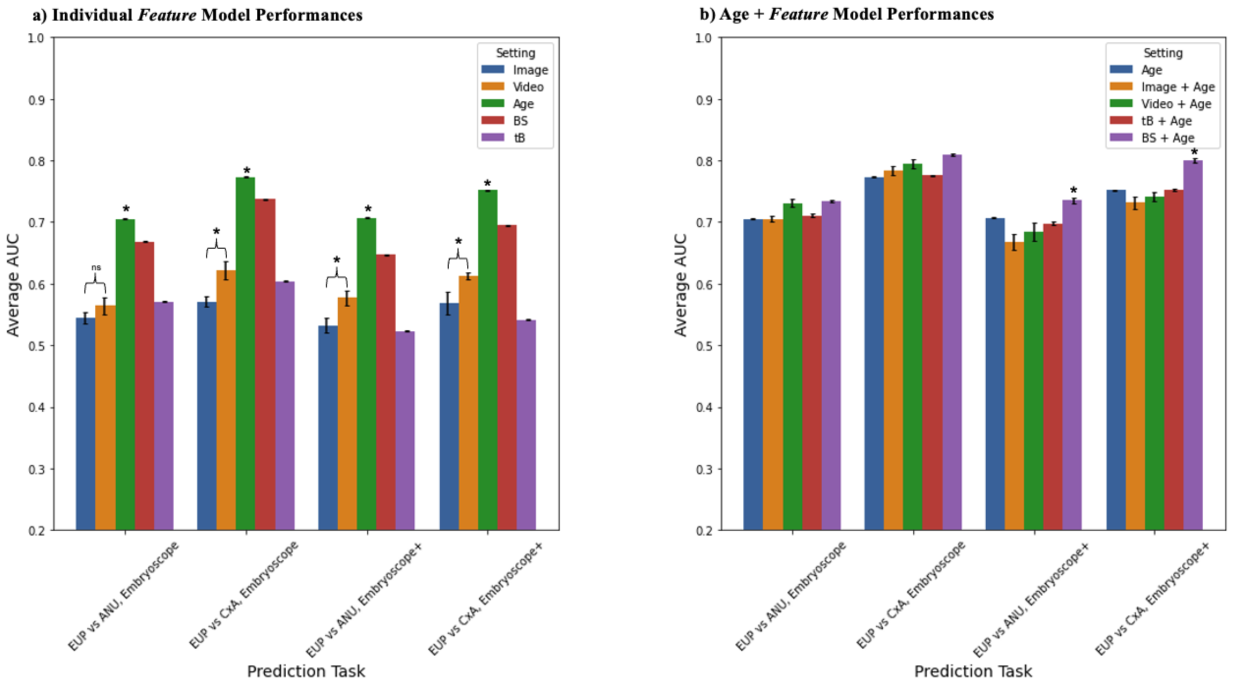


**Supplemental Figure 10. Performance of models with different features across all settings.** Average AUC with standard errors is shown for all prediction tasks on both WCM-Embryoscope and WCM-Embryoscope+ test datasets. (a) Models with only one feature type (image, video, age, blastocyst score, or time to blastulation). (b) Performance of all models with age as a feature.

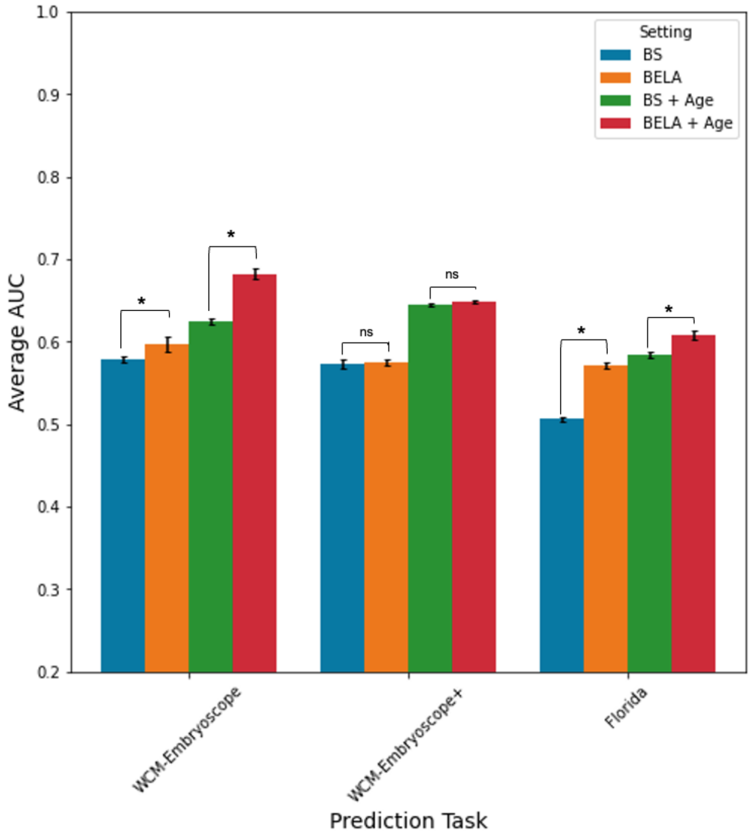


**Supplemental Figure 11. Model performances for Euploid vs Single Aneuploid on all test sets.** Average AUC with standard errors is shown for Euploid vs. Single Aneuploid on all tests using both BELA and embryologist-annotation models. Blue and orange bars show performances of embryologist BS and BELA respectively, without maternal age. Green and red bars show performances of embryologist BS and BELA respectively, with maternal age.

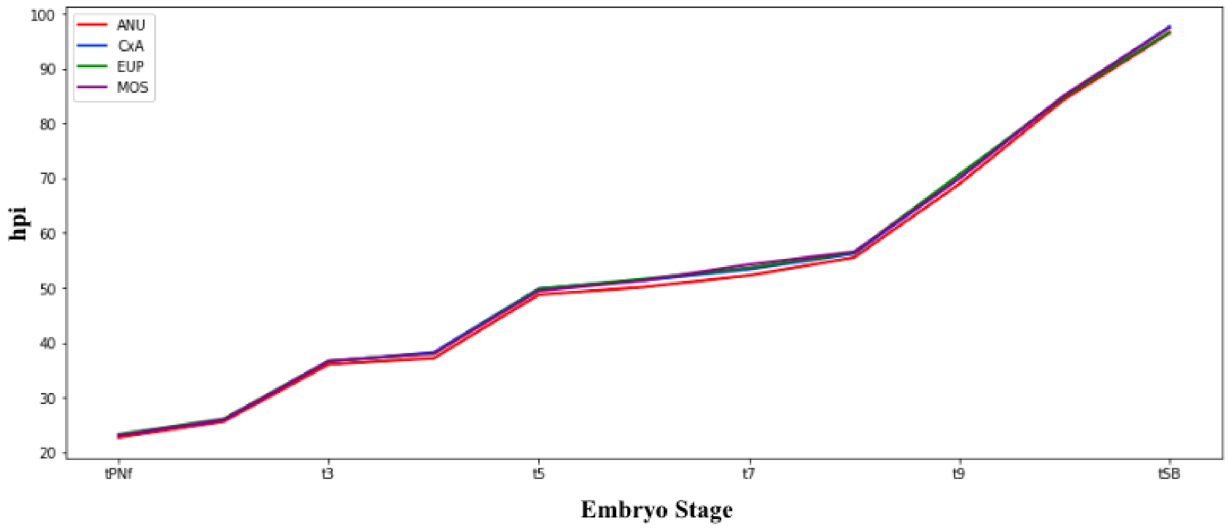

**Supplemental Figure 12. Embryo growth patterns across ploidy via morphokinetic parameters.** Trajectories were extracted from the WCM-Embryoscope+ dataset. X-axis is the embryo stage and y-axis is the hpi.

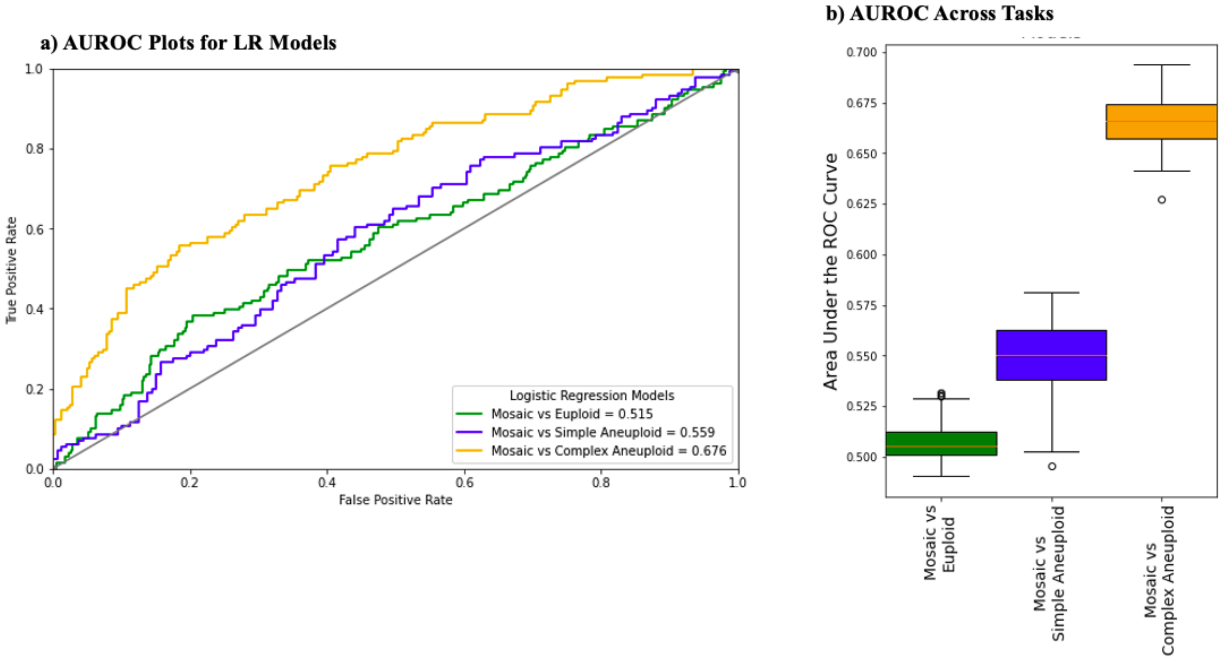


**Supplemental Figure 13. Performances of Logistic Regression models on discriminating mosaic from other ploidy.** (a) shows AUROC plots of mosaic vs euploid, mosaic vs single aneuploid, and mosaic vs complex aneuploid tasks. (b) shows AUROCs across 100 unique iterations of logistic regression training.

#### S3. Supplemental Text

##### Supplemental Text 1. Image versus Video and Time Point Comparison Analyses

To explore the effect of data modalities and focus on specific timepoints for ploidy prediction, we conducted analyses comparing various heterogeneously trained models. Different time point cutoffs were used to restrict the video and image within the day for the prediction model training. Details and justifications of the parameters and training data for each prediction model are provided in **Supplemental Table 3**. For time point and data type comparison analysis, a selection of frame(s) was extracted from the time-lapse sequences of each embryo, depending on the prediction task, as described in the Temporal and Spatial Processing section. Information about selection criteria and parameters for each task are in **Supplemental Table 3**. Embryos with missing clinical information were removed from the dataset. The WCM-Embryoscope data served as our training dataset, reserving 30% for internal testing.

We evaluated prediction model performance using two tasks: EUP versus ANU and EUP versus CxA. Time-lapse sequence images underwent spatial and temporal preprocessing to minimize bias as shown in **Figure 1**, steps 1 - 3 (also refer to Methods). Feature vectors were extracted from processed time-lapse sequences using a pre-trained VGG16 convolutional neural network (**Figure 1**, step 4). We applied various deep learning architectures to predict ploidy from these feature vectors and assessed model performances across time periods and tasks. For video classification tasks, a Bidirectional Long Short-Term Memory network (BiLSTM) architecture was used whereas for image classification tasks, eXtreme gradient boosting tree (XGBoost) model was used (**Supplemental Figure 5**). The BiLSTM received 512-dimensional feature vectors extracted per frame for each embryo whereas the XGBoost accepts a 512-dimensional feature vector from each embryo, corresponding to the chosen image's timepoint. Performance comparison of XGBoost with other models such as logistic regression, random forest, and support vector machines revealed that it either matched or significantly outperformed these alternatives across all tasks. Furthermore, XGBoost surpassed a deep convolutional neural network trained end-to-end, constructed atop a VGG-16 pre-trained model. Maternal age was incorporated as a variable in certain prediction tasks as a continuous variable. Models relying solely on clinical features (embryologist-derived BS, maternal age) utilized a logistic regression architecture, outputting a binary classification (either EUP or ANU/CxA, depending on the task). Four-fold cross-validation was employed to train all models, resulting in four models per task. Model performance was evaluated using accuracy, AUC, precision, and recall on both the WCM-Embryoscope test set and the WCM-Embryoscope+ dataset. Performances were averaged across the four cross-fold models.

**Time point comparison analysis reveals critical timepoints for predictive models**

**Supplemental Figure 6** presents the performance of video and image classification models at different time periods on the WCM-Embryoscope test set. AUC values for video-only classification tasks (**Supplemental Figure 6c,d**) ranged from 0.5 to 0.622, while those for image-only tasks (**Supplemental Figure 6a,b**) ranged from 0.507 to 0.571. For both video-only and image-only classification tasks, model performance on day 5 was significantly higher than on any other day ($p < 0.05$). The performance on days 1-4 showed no statistically significant differences from one another for both image-only and video-only classification tasks. AUC values for video+age classification tasks ranged from 0.719 to 0.795, whereas those for image+age tasks ranged from 0.682 to 0.793 (**Supplemental Table 4**). For both image and video classification tasks that incorporated maternal age, there were no significant differences in performance across time periods.

**Supplemental Figure 7** displays the performance of all non-age models across different time periods on both the WCM-Embryoscope test set and the WCM-Embryoscope+ dataset. For all prediction tasks, day-5 models significantly outperformed other day models. In both video and image classification tasks, there were no statistically significant differences in performance on days 1-4 for the WCM-Embryoscope dataset. The performances across time periods for video+age and image+age models on the WCM-Embryoscope+ dataset are presented in **Supplemental Table 5**. Interestingly, no significant differences between time periods were observed when maternal age was included as an input, suggesting that maternal age reduces the benefits of selecting specific time periods.

To further investigate key time periods for discriminating embryo ploidy status, we conducted principal component analysis (PCA) on the CNN-extracted spatial features for all embryos within the WCM-Embryoscope dataset. Principal components (PCs) were calculated at each time point using the CNN-extracted features derived from the corresponding time-lapse frame. Around 95% of the variance in the extracted features was captured by 100 principal components. Data points were segregated by ploidy status, and inter-cluster distances within a 100-dimensional subspace were calculated for all time periods, measuring from one cluster's center to another. **Supplemental Figure 8** illustrates the intercluster differences at various time periods for both the EUP versus ANU and EUP versus CxA splits. Intercluster differences in both splits increased rapidly after 100 hpi, which occurs on day-5. This finding aligns with the superior performance of day-5 models in discriminating ploidy status. Interestingly, there was a slight increase in intercluster difference between 42 and 46 hpi. This increase, however, was not reflected in day-2 model performances and may be negligible and caused due to random fluctuations. As expected, intercluster differences in the EUP versus CxA split were generally higher than in the EUP versus ANU split, likely due to the more extreme nature of complex aneuploid embryos. In summary, our results indicate that the day-5 time period contains significantly more relevant information for predicting ploidy status compared with other time periods.

**Image and video comparison analysis reveals importance of temporal information**

**Supplemental Figure 9** presents the performance of all day-5 models on both the WCM-Embryoscope and WCM-Embryoscope+ datasets. day-5 was selected as the primary time point for image/video comparison, as our time point analysis indicated that day-5 provided valuable information for ploidy discrimination. In the WCM-Embryoscope dataset, video classification models (video-only and video+age) significantly outperformed image classification models in all prediction tasks ($p < 0.05$). Video-only models also performed significantly better than image-only models in all tasks on the WCM-Embryoscope+ data ($p < 0.05$). Video+age and image+age models exhibited comparable performance on the WCM-Embryoscope+ dataset. Comparisons of image and video classification models on days 1-4 can be found in **Supplemental Tables 4 and 5**. No significant differences were observed between video-only and age-only models at earlier time periods. Video+age models significantly outperformed image+age models on both the WCM-Embryoscope and WCM-Embryoscope+ datasets across days 1-4.

Video-only and image-only models performed significantly worse ($p < 0.05$) than models trained solely on blastocyst score (BS) or maternal age (**Supplemental Figure 10a**). The BS is a quality metric based on the conversion of embryologist-annotated morphological grades for blastocysts on day-5. Time to blastulation (tB)-only models exhibited comparable performance to video-only classification models on the WCM-Embryoscope dataset. Although video+age and image+age models performed significantly better than age-only models on the WCM-Embryoscope dataset, they performed significantly worse on the WCM-Embryoscope+ data (**Supplemental Figure 10b**). These results collectively suggest that including maternal age reduces the distinction between image and video performance, particularly when models are tested on a heterogeneous dataset like the WCM-Embryoscope+. In general, however, video-based models significantly outperformed single-image-based models, likely due to the added temporal information.

Our results suggest that day 5 of embryo development provides the most information for ploidy discrimination. Day-5 models also perform significantly better than all-day models as well (**Supplemental Table 4**) suggesting that information, or lack thereof, detracts from model performance. Our second analysis compared day-5 image classification models to video classification models. Video-only models performed better than image-only models at all prediction tasks on both the WCM-Embryoscope and WCM-Embryoscope+ datasets. Video+age models also performed significantly better than image+age models in all prediction tasks on WCM-Embryoscope data. These results suggest that the temporal information from additional frames provides models with heightened ability to classify embryos by their ploidy status. That said, embryologist-derived quality metrics within the Weill Cornell datasets like blastocyst score are significantly more predictive of ploidy status compared with both image and video data. This suggests that video and image classification models are unable to encapsulate different aspects of embryo development that embryologists use to determine the blastocyst score. Features extracted from days 1-4 may not be informative and could decrease overall model performance in all-day models, as compared to day-5 models. This is consistent with the biological nature of embryo development, as specific features associated with chromosomally normal embryos, such as cavitation, occur on day 5.[10] Other studies have suggested that other morphological characteristics of the embryo that occur early in development may be suggestive of an embryo's potential to implant.[22] These features include those such as rate of cleavage, cell count, and length of morula stage.[22] However, many of these studies have different definitions of what a "successful" embryo is, varying from an embryo that reaches the blastocyst stage to an embryo that leads to a pregnancy. Although a chromosomally normal embryo may be correlated with these success criteria, the indicators may not necessarily be the same. Our study suggests that these early indicators of embryo "success" are not suggestive of ploidy status. We hypothesize that the rate of cavitation and speed to blastulation may be markers for ploidy that our video classification models are able to extract (**Supplemental Figure 10a**). Interestingly, when maternal age was added as a feature to the models, there were no significant differences across time periods (**Supplemental Figure 6**). Performance across all prediction tasks and datasets was higher with the addition of maternal age. This suggests that maternal age is a strong predictor of ploidy status, which is consistent with previous studies, and stronger than both video and image data (**Supplemental Figure 10a**). As far as we know, this is the first study that comprehensively investigated modality and temporal information within time-lapse imaging. Based on these results, we suggest that future efforts to discriminate embryos on ploidy status focus on later stages of embryo development.

#### Supplemental Text 2. BELA Model for Euploid versus Single Aneuploid Discrimination

To further elucidate the predictive capabilities of the BELA model, a focused experiment was conducted, differentiating euploid embryos from single aneuploid embryos. Single aneuploid embryos are significant, as they are frequently associated with miscarriages and abnormal offspring outcomes. In contrast, complex aneuploidies are more commonly tied to outcomes like embryo arrest or implantation failure.

For this investigation, we formulated four distinct models. The first was the BELA model, which utilized embryonic time-lapse data without the inclusion of maternal age. In the second model, we incorporated both the embryonic time-lapse data and maternal age into the BELA framework. The third model was constructed solely based on the manually annotated embryologist evaluations of the blastocyst score. Finally, the fourth model combined both embryologist annotations and maternal age as determining factors. Each of these models adhered to the architectural designs and training protocols detailed in the Methods section of the main manuscript. By comparing the performance of these models, we aimed to validate the efficacy of the BELA model while also gauging its effectiveness relative to models relying on traditional and biologically pertinent data. To validate the performance of the models, they were evaluated on the WCM-Embryoscope, WCM-Embryoscope+, and Florida test sets (**Supplemental Figure 11**, **Supplemental Table 11)**. In this analysis, the Spain dataset was removed as it did not provide any discrimination on whether an embryo was single or complex aneuploid.

Both BELA models – with and without maternal age – consistently outperformed their embryologist-annotated blastocyst score (BS) counterparts in the WCM-Embryoscope and Florida datasets. However, a notable exception was observed in the WCM-Embryoscope+ dataset where the performance of the BELA model was found to be on par with the embryologist-annotated BS models. It is important to highlight that the performance metrics for all models in the EUP vs. SA task were generally lower than when distinguishing euploid from any type of aneuploid embryo. For instance, in the WCM-Embryoscope dataset, the performance of the BELA model with maternal age in the EUP vs. ANU (any aneuploidy) task rendered a value of 0.76 ± 0.002. However, when its counterpart EUP vs SA had a performance of 0.682 ± 0.005 ($p < 0.05$). An intriguing observation was made in the Florida dataset. Here, the embryologist annotated BS appeared to offer no predictive advantage when discriminating between EUP and SA. Yet, even the BELA model without the inclusion of maternal age demonstrated significantly improved performance. This superior efficacy of the BELA model might be attributed to the enhanced granularity present in the Weill Cornell grading system. From a biological standpoint, these findings align with our understanding of embryo development. Single aneuploidies inherently pose greater challenges for discrimination compared to their complex counterparts. This is substantiated by instances where embryos with single aneuploidies not only successfully implant but also culminate in viable pregnancies.[23,24]

#### Supplemental Text 3. Investigation of Mosaic Embryos

Within the WCM-Embryoscope+ dataset, 142 embryos were identified as mosaic via PGT-A. Though these were excluded from the primary evaluation of the BELA model, we undertook a comprehensive analysis to discern any distinctions between mosaic embryos and those labeled as euploid or aneuploid. **Supplemental Figure 12** presents a comparative assessment of embryo growth trajectories across varied ploidy classifications, referencing specific morphokinetic parameters. This illustration, derived from the WCM-Embryoscope+ dataset, delineates the mean growth trend aligned with each ploidy status. Intriguingly, the growth patterns across these statuses exhibit marked congruence, underscoring a minimal divergence. Consequently, based on the evaluated morphokinetic parameters, distinguishing between mosaic embryos and those of alternate ploidy statuses remains challenging.

To ascertain the unique characteristics of mosaic embryos, logistic regression classifiers were constructed with three distinct aims: (1) differentiate between mosaic and euploid embryos, (2) differentiate between mosaic and single aneuploid embryos, and (3) differentiate between mosaic and complex aneuploid embryos. The training was conducted on the WCM-Embryoscope+ dataset, utilizing maternal age, morphokinetic parameters, and manual embryologist quality scores as features. Given the dataset's constraints, the area under the receiver operating characteristic curve (AUROC) was determined using the classifier's out-of-bag (OOB) predictions. This prediction methodology was chosen to ensure model generalizability, mitigate overfitting, and permit training utilizing the entire dataset. **Supplemental Figure 13a** illustrates the AUROC plots of the OOB predictions. Concurrently, **Supplemental Figure 13b** presents a box and whisker diagram delineating the area under the curve (AUC) for each classifier over 100 distinct training iterations. The mosaic versus euploid classifier demonstrated results near random levels for mosaic embryo differentiation. While the mosaic vs single aneuploid classifier exhibited marginally enhanced performance compared to its euploid counterpart in identifying mosaic embryos, its efficacy remained proximate to random levels for this task. Notably, the mosaic vs complex aneuploid classifier displayed a significant advancement in performance. These findings indicate that the chosen features facilitate differentiation between mosaic and complex aneuploid embryos but do not offer the same clarity when distinguishing mosaic embryos from those that are euploid or single aneuploid.

Our evaluations indicate that mosaic embryos bear significant resemblance to euploid embryos. Given these findings, we concluded that excluding mosaic embryos during BELA's training would be optimal due to the restricted sample size of mosaic embryos, notably absent in our WCM-Embryoscope training set. Moreover, the clinical interpretation and PGT-A results of mosaic embryos are still unclear.[25]

#### Supplemental Text 4. Adding Additional Training Data to BELA

In an effort to evaluate the potential benefits of training BELA on heterogeneous data, we retrained the model using a combination of the initial WCM-Embryoscope and Florida datasets. However, due to differences in the blastocyst scoring methods between the Weill Cornell and Florida datasets, we deemed it unsuitable to train the first component of BELA (blastocyst score prediction module) on the Florida dataset. Consequently, the first component of BELA was trained in the same manner as detailed in the main text. For the second component, we integrated information from the Florida dataset, predicting MDBS for both Weill Cornell and Florida data, and subsequently feeding it into a logistic regression model to predict ploidy status. The retrained BELA did not exhibit significant improvements compared with the original BELA presented in the main text across any analysis. Nonetheless, several limitations should be noted. Firstly, the inability to fully utilize the Florida dataset (by training the blastocyst score prediction module) was due to the differences in blastocyst ground truth derivation between the two sites. Secondly, advanced video classification architectures, such as 3D-CNNs or Inflated 3D ConvNets—which exceed the scope of this manuscript—might be more adept at capturing dataset heterogeneity, potentially leading to enhanced results. These aspects will be the focus of future investigations.
